# Emergence of Genetic Sex Determination in an Environmentally Sex-Determined Animal

**DOI:** 10.1101/2025.09.08.674940

**Authors:** Wen Wei, Man Lin, Trent Bishop, Ryan Shahriari, Yue Hao, Logan Graham, Bret L. Coggins, Hannah Lew, Swatantra Neupane, Li Wang, Sen Xu, Erin Kim, Michael Lynch, Zhiqiang Ye

## Abstract

The evolution of genetic sex determination (GSD) from environmental sex determination (ESD) remains a fundamental issue in evolutionary biology. However, the mechanisms driving such a transition, particularly in its earliest stages, are largely unknown. Here, we report on a genetic variant in the ESD species, *Daphnia pulex*, in which structural variations and selection on chromosome Ⅰ have suppressed recombination. This suppression has facilitated the accumulation of antagonistic features of the two chromosome-Ⅰ haplotypes, resulting in heterozygosity and differentiation of the sex-determining gene *Dfh* on two haplotypes. Consequently, this process has driven the emergence of a mixed-mating system, in which sex determination is jointly regulated by genetic and environmental factors. Additionally, we detected ongoing divergence in chromosomal structure between the two haplotypes at the population level. These findings suggest that processes analogous to early sex-chromosome evolution are occurring in this ESD species, offering novel insights into the molecular mechanisms underlying the emergence of GSD.

## Introduction

Many fundamental questions in evolutionary biology center on how reproductive system diversity arises and is maintained across diverse phylogenetic lineages ^1–3^. In animals, the evolution of sexual dimorphism and the emergence of sex chromosomes, which establish alternative mating systems, represent fundamental aspects of biology. These processes often involve transitions between two major modes of sex determination ^4,5^: environmental sex determination (ESD), influenced by factors such as temperature and photoperiod; and genetic sex determination (GSD), controlled by sex chromosomes or chromosomal regions.

Theoretical models propose several key steps in the evolutionary transition from ESD to GSD ^6–9^. First, an upstream mutation that causes sterility in one sex—typically male—results in a female-determining allele and the emergence of gynodioecy, a breeding system in which genetic females coexist with hermaphrodites ^10–12^. These sex-determining mutations may arise in transcription factors such as *sisterless* in fruit flies ^7,10^ and *sry* in mammals ^11–13^, from the TGF-β signaling pathway (e.g., *amhy* in teleost fishes ^14,15^), or from a hormone-related pathway ^16^. Gynodioecy is widely observed in plants and is considered an intermediate stage in the evolution of sex chromosomes, but it is extremely rare in animals ^9,17–21^. Second, recombination suppression between emerging proto-sex chromosomes evolves through mechanisms such as sexually antagonistic selection ^21–23^, a ‘lucky’ inversion ^24^, or the neutral accumulation of variants near the sex-determining genes via genetic drift ^25–27^. Third, as sexually antagonistic genes—whose functions favor either sex—accumulate near the sex-determining region, the recombination-suppression region of the proto-sex chromosomes expands ^9,28–32^. Finally, one of the proto-sex chromosomes degenerates over time ^28,29^, potentially leading to the heteromorphic structures that characterize fully differentiated sex chromosomes.

While extensive research has focused on the downstream divergence of proto-sex chromosomes, relatively few studies have investigated the early transitions, particularly the shift from ESD to gynodioecy ^33–36^. In this study, we focus on a recently discovered variant in the reproductive system of the microcrustacean *Daphnia pulex*, a species that predominantly relies on ESD ^30^. *Daphnia* typically reproduce via cyclical parthenogenesis, where females produce daughters asexually for an indefinite number of generations. However, environmental changes can trigger shifts in the juvenile hormone (JH) and ecdysone (20E) pathways, leading to the production of males ^31,33^. Recent findings in *D. pulex* reveal the existence of loss-of-ESD genotypes that are incapable of producing males under male-induction cues (Figure 1a), a trait apparently controlled by a major dominant variant at a single segregating locus ^34^. The coexistence of non-male-producing (NMP) clones (genetically determined females) and ESD clones resembles gynodioecy, providing a rare and potentially powerful model to explore the early steps of sex chromosome evolution.

**Figure 1:**
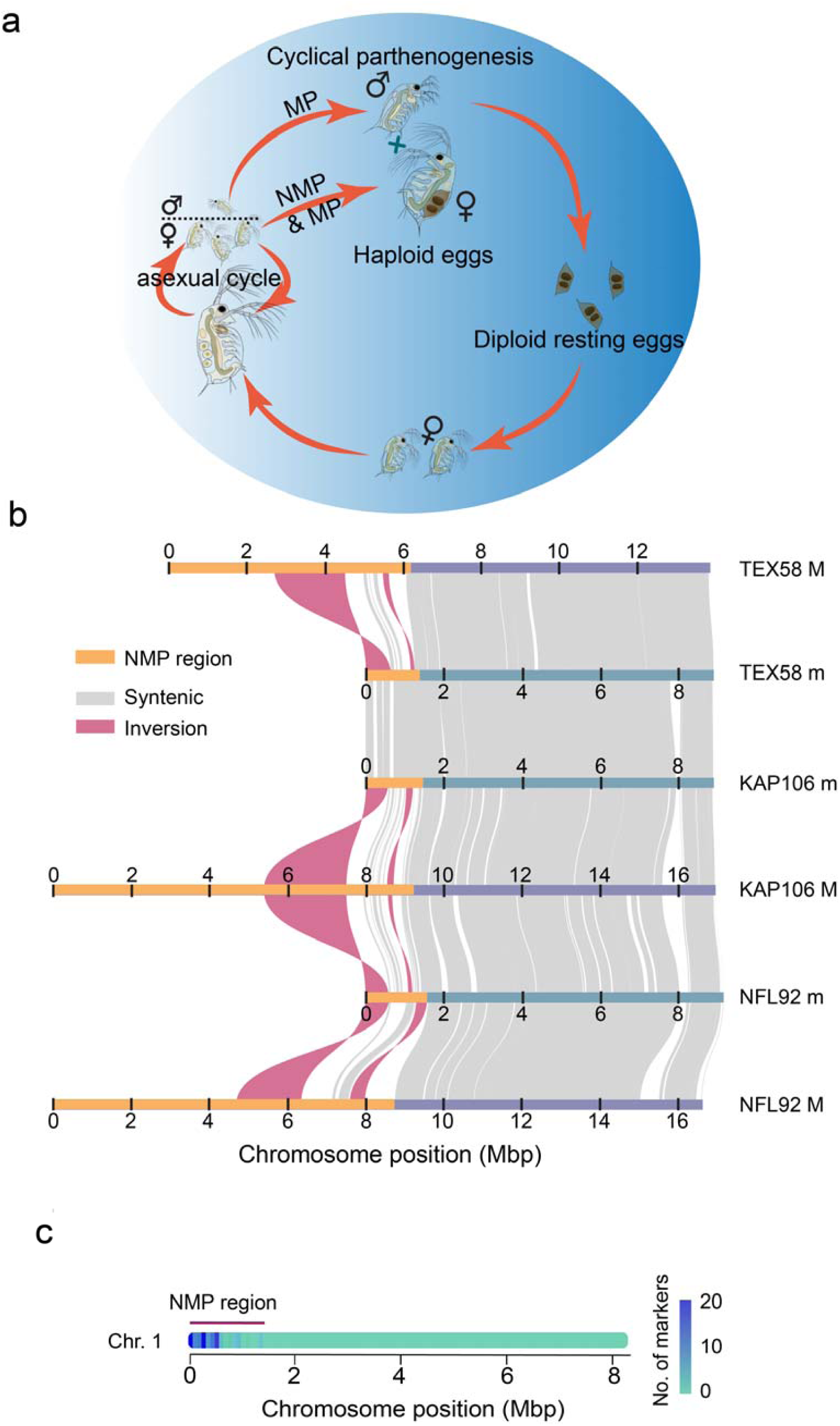
Heteromorphic chromosomes in NMP *D. pulex*. **a)** Reproductive modes in *Daphnia*. *Daphnia* exhibits two reproductive strategies. MP (male-producing) genotypes generate males and females, whereas NMP (non-male-producing) genotypes only produce female offspring. In both cases, females can continue to reproduce clonally for extended periods. Sexual reproduction produces diploid diapausing resting eggs, which eventually hatch into female offspring. **b)** Syntenic comparison of haplotypes from three NMP clones, using each *m* respective haplotype as the mapping reference. This analysis demonstrates that the *m* haplotypes of all isolates closely resemble each other, whereas the *M* haplotype restricted to NMP clones exhibits substantial structural variation among populations. Syntenic regions and inversions more than 20 kb are displayed. **c)** Distribution of NMP markers in Chromosome I based on the KAP4 reference genome. The colored block in the panel represents the number of NMP markers within 1-kb windows.

To investigate the mechanisms underlying this transition, we characterized the sex-determining region (SDR) in *D. pulex* and identified *Dfh* as a key regulator of sex determination. Moreover, we analyzed the genomic features that lead to recombination suppression within this region, offering new insights into how genetic mechanisms can override environmental cues. Together, these findings enhance our understanding of the molecular basis for the evolutionary transition from ESD to GSD.

## Results

### Identification and Comparative Analysis of the NMP Region

Because *Daphnia pulex* is ancestrally capable of producing males, the non-male-producing (NMP) phenotype must have arisen from variants within a male-producing (MP) genetic background. Previous work implies that the NMP trait is controlled by dominant heterozygous loci, likely linked to chromosome I ^34^. However, the fragmented assembly of an earlier version of the *D. pulex* reference genome limited the precision of NMP-associated loci mapping, and the lack of a reference genome from an NMP clone has impeded efforts to investigate structural variation (SV) potentially underlying the trait.

To pinpoint the genomic basis of this heterozygous region, we assembled a chromosome-scale genome for a North American MP clone (KAP4; Figure S1a-b), using 108× PacBio HiFi and 191× Hi-C data. The resulting assembly spans 133.2 Mb across 12 chromosomes, each resolved into one or two contigs—marking a significant improvement over the earlier draft ^37,38^. With a BUSCO completeness score of 98.1% (Figure S1b), the KAP4 reference genome supports the annotation of 22,355 genes, including 15,282 protein-coding and 7,073 non-coding genes. In parallel, we generated fully phased diploid genome assemblies for three NMP clones obtained from isolated populations (KAP106, NFL92, and TEX58; Figure S1a-b), using PacBio HiFi coverage ranging from 113× to 254× and ∼200× Hi-C. Each haplotype was assembled into ∼20 contigs and confidently anchored to the 12 chromosomes. The haplotypes of the NMP clones are comparable in size to the KAP4 reference genome across most chromosomes (Figure S1c). However, a striking difference is observed on chromosome I within NMP clones: one haplotype closely matches the MP genome (∼9.0 Mb), while the other is greatly expanded and variable in size (17.0 Mb in KAP106, 16.6 Mb in NFL92, and 13.8 Mb in TEX58), suggesting that this chromosome harbors the determinants of the NMP phenotype.

We conducted structural genome comparisons among the NMP clones and KAP4 to further investigate the basis of length variation on chromosome I (Figure 1b; Figure S1d). The shorter chromosome I haplotypes from the NMP clones display near-complete collinearity across clones. In contrast, the longer haplotypes exhibit substantial SV localized to the proximal region of the chromosome, spanning 0–9.2 Mb in KAP106, 0–8.8 Mb in NFL92, and 0–6.2 Mb in TEX58, while retaining collinearity across the remaining length of the chromosome (Figure 1b). The corresponding region is located at position 0–1.4 Mb on KAP4 chromosome I and in the shorter chromosome I haplotypes in NMP clones. The NMP phenotype is associated with a heterozygous linkage haplotype (designated as *Mm*), while MP clones are homozygous (*mm*). This divergent NMP region of the *M* haplotype harbors SVs that are absent in the corresponding *m* haplotype. In addition, all *M* haplotypes share a large-scale inversion spanning 0.06–0.60 Mb relative to the KAP4 reference. This inversion extends to approximately 2 Mb in the NMP clones, spanning 5.4–7.5 Mb in KAP106, 4.3–6.4 Mb in NFL92, and 2.6–4.5 Mb in TEX58. This divergent structural rearrangement, confined to NMP chromosomes, suggests that the inversion predates the divergence of the NMP haplotype lineages.

To investigate the underlying causes of the pronounced size expansion observed in the *M* haplotype, we evaluated two potential mechanisms: 1) an increase in gene content, due to duplication or translocation; and 2) the accumulation of repetitive, non-coding DNA. First, comparative gene annotation revealed that candidate NMP region of the *M* haplotype harbors a substantially elevated number of protein-coding genes, averaging 806 per haplotype, in contrast to only 115 in the corresponding region of the *m* haplotype. Across the *M* haplotype, 217 genes are shared among all three variants (Table S1a), with the remainder being unique to individual clones. Of the genes present in the *m* haplotype, 103 are also found in the *M* haplotype. Notably, eight of 103 have undergone copy number expansion in the *M* haplotype, ranging from two to eleven copies, whereas their *m* haplotype counterparts remain single copy. After excluding these shared genes, we identified 96 *M*-specific genes. These genes originated from interchromosomal duplications involving regions outside the NMP locus, including non-NMP segments of chromosome I as well as chromosomes II through XII (Figure S1e).

To estimate the timing of these duplication events, we applied the molecular clock equation, *T = ΔS / 2μ*, where *μ=*4.33 × 10⁻⁹ substitutions per site per generation, as previously inferred for a cyclical parthenogenic *D. pulex* ^39^, and *ΔS* represents synonymous divergence between gene copies in the *M* haplotype vs. the *m* haplotype (specifically from KAP4). We averaged the estimated pairwise ages of multi-copy genes, as complex duplication histories make it difficult to distinguish individual duplication events and identify the original copy. The median estimated divergence time between duplicated genes and their ancestral copies is approximately 0.97 (SE = 0.14) million years ago (MYA) (Figure S1f). This estimated divergence time predates the inferred origin of the NMP haplotype (0.19–0.85 MYA ^34^), which was based on a phylogenetic analysis of gene 8960, a potential contributor to the NMP phenotype.

Second, we observed a marked enrichment of repetitive elements within the *M* haplotypes, primarily owing to retrotransposons, DNA transposons, and fragmented mobile elements (Table S1b). Repeats constitute, on average, 38.2% of the *M* haplotype, significantly exceeding the 9.7% observed in the *m* haplotype across three independent NMP clones (Mann–Whitney U test, *P* = 0.029). This discrepancy is particularly pronounced within the inversion region associated with the NMP haplotype, where repeat content reaches 70.5% in *M* versus only 8.0% in *m* (*P* = 0.025). In contrast, no significant difference in repeat abundance is detected outside the NMP region of chromosome I (9.6% in *M* vs. 9.0% in *m*; *P* > 0.05), suggesting that repeat accumulation is localized and not genome-wide.

To assess whether such repetitive sequences are shared across *M* haplotypes, we examined the 217 genes common to all three *M* haplotypes, along with their flanking regions. Most TEs are not located near these genes; for example, in KAP106, only 70 out of 217 genes contain or flank TEs. Due to extensive population-specific rearrangements, we excluded distal intergenic regions from this analysis, as alignment of homologous sequence is not feasible. Among the shared loci, only seven retrotransposons, located within or flanking six genes, are conserved across haplotypes. The limited overlap in both duplicated gene content and repeat elements implies that the bulk of structural expansion in *M* haplotypes represents lineage-specific events. These patterns suggest that most of the observed genomic expansion is unlikely to be causally associated with the origin of the NMP phenotype, but rather reflects independent trajectories of structural genome evolution.

Given that the NMP phenotype results from a loss of male-producing function, we sought to identify shared signatures of potential loss-of-function alleles across *M* haplotypes. Specifically, we focused on two forms of disruption: 1) complete gene loss; and 2) mutation accumulation that may impair gene function. First, we found 12 genes that are consistently absent from all *M* haplotypes when compared to the MP reference genome (KAP4) (Table S1c). Second, to detect shared sequence-level disruptions, we analyzed single-nucleotide variants (SNVs) by mapping genomic reads from 100 MP and 67 NMP clones (from our previous study ^34^) to the KAP4 reference (Table S1d). This analysis again pinpointed the NMP-associated region to the 0–1.4 Mb segment, where we phased 427 SNVs (Table S1e) that reliably distinguish the *M* haplotype from the *m* haplotype (Figure 1c). These SNVs span 28 candidate genes for functional divergence. Notably, 84.5% of these NMP-associated SNVs fall within the ancestral inversion, further implicating this structural variant as having a role in recombination suppression and potentially facilitating the accumulation of mutations contributing to the NMP phenotype. Together, the 12 genes absent from *M* haplotypes and the 28 genes harboring NMP-associated SNVs could be potential candidates to contribute to the loss-of-ESD.

### Identification and Characterization of the Sex-Determining Gene Dfh

In *Daphnia*, the production of male offspring is typically triggered by environmental cues perceived by the mother, such as photoperiod or population density ^40–45^. These environmental cues are transduced through maternal hormonal signals that act on developing oocytes ^45,46^. Because we did not observe embryo lethality in the brood chambers of NMP mothers, we hypothesize that the NMP phenotype likely arises not from male-specific lethality, but from genetic variation that blocks male determination. This could result from allelic variants that disrupt the male-determination pathway in offspring (embryos), thereby biasing development toward the female trajectory, or that impair maternal transmission of environmental cues required to initiate male determination. To evaluate these possibilities, we analyzed the candidate genes at both the embryo and maternal level, specifically the 12 genes absent from *M* haplotypes and the 28 genes harboring NMP-associated SNVs (see above mentioned), to assess their potential roles in the origin of the NMP trait.

We first identified genes that are differentially expressed between male and female embryos during the development window in the brood chamber (∼3 days). To this end, we performed RNA-seq analyses across multiple key developmental stages, including 12-h, 36-h, and 60-h embryos (E-12 h, E-36 h and E-60 h), under two environmental conditions known to bias sex determination (Figure 2a; Table S2a). We compared embryos reared under short-day (SD) photoperiods (10 h light/14 h dark), which induce male production in MP clones but not in NMP clones, and long-day (LD) photoperiods (14 h light/10 h dark), which result in female-only offspring in both genotypes (Figure S2a).

**Figure 2:**
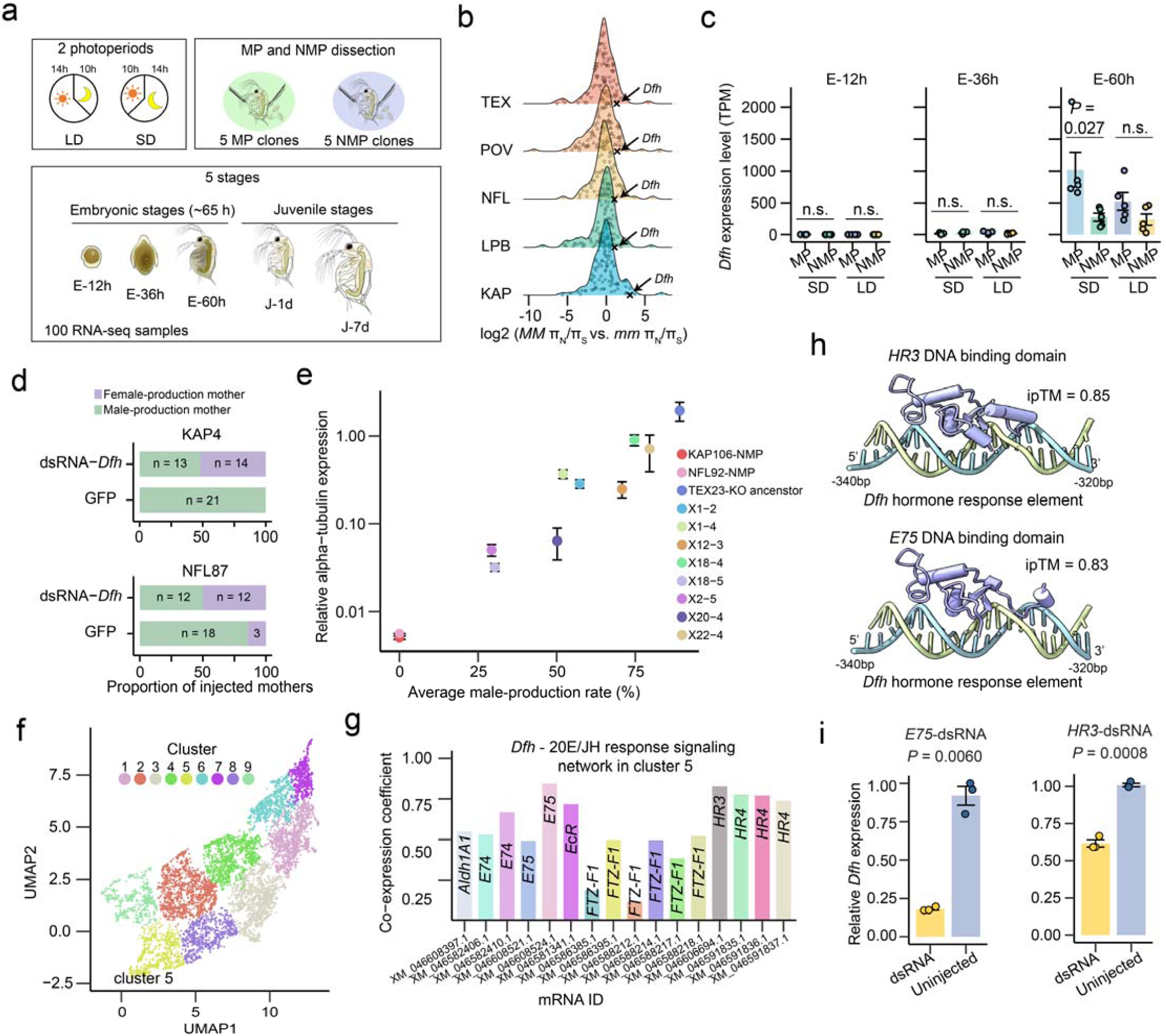
Hormonal regulation of male-determining gene *Dfh* expression. **a)** RNA-seq sampling workflow. MP clones (NFL87, TEX23, LPB120, POV15, KAP4) and NMP clones (NFL92, WVA79, KAP106, BRG89, TEX58) were sampled at five developmental stages: 12-hour (E-12h; early stage), 36-hour (E-36h), and 60-hour (E-60h; late stage) embryos, as well as one-day (J-1d) and seven-day (J-7d) juveniles, under long-day (LD) and short-day (SD) conditions. Embryos were dissected from one-month-old mothers for collection. **b)** Distribution of the *MM* π_N_/π_S_ ratio relative to the *mm* π_N_/π_S_ ratio for NMP region genes. Only common genes between *M* and *m* haplotypes were calculated. The fold change in the π_N_/π_S_ ratio of *MM* relative to *mm* is plotted for genes. Each dot represents a gene. **c)** *Dfh* expression levels in MP and NMP clones. The TPM (Transcripts Per kilobase Million) values of *Dfh* are shown for each clone. Each dot represents a clone, and the data are presented as mean ± S.E.M. (standard error of the mean). *P*-values were obtained using *t*-tests comparing MP and NMP clones. **d)** Knockdown of *Dfh* affects male production. The number of male-producing mothers was compared between the *Dfh* knockdown group and the control group (injected with GFP). Data from two MP clones (KAP4 and NFL87) show a reduced number of male-producing mothers in the *Dfh* knockdown group. **e)** Relationship between *Dfh* expression and male-production rate in CRISPR knockout lines and the ancestral TEX23 clone under methyl farnesoate (MF) conditions. Individuals were exposed to MF for at least two weeks, and 60-hour embryos from >15 individuals per clone were collected for RNA extraction. Expression levels are reported as mean ± S.E.M. from three primer sets spanning exons 1–2, 2–3, and 3–4, each based on the mean of three qPCR technical replicates. **f)** Clustering of differentially expressed genes (DEGs). Using the UMAP algorithm, DEGs in SD samples were clustered based on their TPM values, resulting in 9 distinct expression clusters, indicating unique transcriptional patterns. **g)** Correlation coefficients of TPM values across SD samples between *Dfh* and genes involved in the juvenile hormone (JH) and ecdysteroid (20E) signaling pathways within Cluster 5. Cluster 5, which includes *Dfh*, contains key genes involved in JH and 20E signaling. The x-axis represents mRNA IDs, corresponding to alternative splice isoforms or duplicated gene copies for these genes. **h)** Predicted interaction between *Dfh* regulatory region and DNA-binding domains of *E75* and *HR3*. We analyzed the 1 kb region upstream of the *Dfh* transcription start site (TSS) for potential binding sites of *E75* and *HR3* and predicted interactions using AlphaFold3. The structure features helices (cylinders), sheets (arrows), and DNA, with predicted scores (called ipTM values) above 0.8 indicating highly confident predictions of protein-DNA interaction. **i)** qPCR validation of dsRNA knockdown effects. RNA was extracted from more than 20 individuals at day three post-injection. *Dfh* expression levels were measured. Each dot represents a qPCR replicate, with data shown as mean ± S.E.M. Statistical significance was assessed using *t*-tests comparing dsRNA-injected and uninjected groups.

Under SD conditions, MP clones produced male embryos, whereas NMP clones exclusively produced females, suggesting that the difference lies in the regulation of sex-determining genes during embryogenesis. To identify candidate genes, we compared gene expression between five MP and five NMP clones under SD conditions across each developmental stage. Because some expression differences may reflect broader genetic divergence unrelated to sex determination, we filtered out genes that were also differentially expressed between MP and NMP clones under LD conditions— where both genotypes produce only females—to focus on genes specifically associated with male production (Figure S2b; Table S2b).

Among the 12 genes absent from the *M* haplotypes, one exhibited significantly reduced expression in NMP clones (via the intact *m* copy) compared to MP clones at the E-36 h stage. The expression was reduced by more than twofold (3.8-fold), suggesting that the decrease cannot be explained solely by the absence of a gene copy (Figure S2c). Additional downregulation is likely caused by other regulatory controls within the NMP region. On the other hand, 4 of the 28 genes harboring NMP-associated SNVs were identified as differentially expressed genes (DEGs) (Figure S2c) and markedly reduced expression of the *M*-haplotype allele relative to the *m*-haplotype allele in NMP clones (Figure S2d). These five genes may collectively form the genetic basis underlying the transition away from ESD (Table S2c).

To evaluate potential differences in selection pressures on NMP region-expressed DEGs between the *m* and the *M* haplotype, we compared πN/πS (ratio of nonsynonymous to synonymous nucleotide diversity) for NMP region alleles. For a given genomic site, the heterozygosity across NMP clones can be viewed as an average of the four possible haplotype pairings: π* = 0.25π_MM_ + 0.5π_Mm_ + 0.25π_mm_, where π_mm_ and π_Mm_ represent the average site-specific heterozygosity among *mm* (MP) clones, and between chromosomes within *Mm* (NMP) clones for the gene. Substitution of the latter allows estimation of πMM, the mean diversity across *M* haplotypes of each gene. Notably, only one NMP-region gene (which is also a DEG)—LOC124196108 (one of the five in Table S2c) —consistently exhibited a >2-fold increase in π_N_/π_S_ of *MM* compared to *mm* across all populations (Figure 2b), suggesting either relaxed purifying selection or potential positive selection of this gene in NMP.

This gene, which lies within the inversion region, has 3.7-fold higher expression in MP clones compared to NMP clones at the 60-hour embryonic stage (E-60 h; Figure 2c), a period consistent with the previously identified sex-determination window of 40– 60 hours post-ovulation in *D. pulex* ^44,46^. Allele-specific expression analysis further revealed markedly reduced expression of the *M*-haplotype allele relative to the *m*-haplotype allele, implicating expression level of LOC124196108 as a potential mechanism of male suppression in NMP clones (Figure S2d). Sequence analysis indicated that LOC124196108 is essentially identical to gene 8960, a previously proposed candidate for the NMP trait identified via phylogenetic analysis, with minor differences likely arising from earlier annotation errors (blastn identity of coding-region = 99.6%). The convergence of evidence from differential expression, signatures of selection, and phylogenetic inference strongly supports the deep involvement of LOC124196108 in the NMP phenotype and, more broadly, in loss of environmental sex determination. In light of its putative role in directing male versus female development, we propose naming this gene *Dfh* (*Daphnia feng huang*), after the Chinese phoenix— where “*feng*” symbolizes the male and “*huang*” the female—reflecting its functional implications in sex differentiation.

To test the functional role of *Dfh* in sex determination, we used RNA interference (RNAi) to reduce *Dfh* expression in early-stage male embryos derived from two MP clones (KAP4 and NFL87). In *Daphnia*, male production is typically initiated by the maternal response to methyl farnesoate (MF), an upstream hormonal signal ^31,33^. Given that *Dfh* expression peaks during the embryonic sex-determination window (around 60 hours post-ovulation) and can be upregulated by maternal MF exposure (Figure S3a), we hypothesized that *Dfh* functions downstream in this signaling cascade. If *Dfh* is essential for environmentally determining the sex, then silencing its expression in embryos produced by MF-treated mothers should suppress male determination and reduce the male-production rate in response to MF.

To further explore this hypothesis, we microinjected male embryos with *Dfh*-dsRNA alongside a GFP-reporter construct (*Dfh*-dsRNA+GFP), while control embryos received only the GFP-reporter. GFP expression and embryonic sex were then evaluated at three and five days post-injection (Figure S3b-c). To minimize batch effects on surviving embryos, we evaluated the proportion of male-producing mothers rather than raw embryo counts. *Dfh* knockdown (to ∼30% of control from the qPCR estimates; Fig. S3c) significantly reduced male production relative to controls (100.0% vs. 48.1% in KAP4; 85.7% vs. 50.0% in NFL87; Figure 2d). Histological analysis using hematoxylin-eosin staining confirmed that *Dfh*-suppressed individuals that developed as females exhibited normal ovarian morphology, indicating that loss of male development did not disrupt female gonadal differentiation (Figure S2d).

Because RNAi typically does not completely eliminate gene expression, we additionally generated eight independent *Dfh* mutant lines using CRISPR-mediated knockout (KO) (Table S2d). Under standard culture conditions (clear water, sufficient food, and low density), these KO lines produced no males but the KO ancestor TEX23 did. To further evaluate the consequences of these mutations on both gene expression and sex determination under artificial male-inducing conditions (800 nM MF), we quantified male-production rates for 7–15 individuals per line for three successive clutches, alongside unedited MP clone as a control. In parallel, we measured *Dfh* expression in 60-hour embryos using qPCR. All CRISPR-derived lines exhibited reductions in both *Dfh* transcript levels and male production, revealing a positive association between gene expression and phenotypic response (Figure 2e; Figure S3e). Two lines in particular, X2-5 and X18-5, exhibited a pronounced loss of *Dfh* expression and a substantial decline in male-production rates (∼30% compared to ∼90% in TEX23). Notably, 4 of 15 X2-5 individuals and 3 of 13 X18-5 individuals failed to produce any males over the two-week testing period. Together with the RNAi results, these findings demonstrate that *Dfh* is a critical sex-determination factor: reduced expression largely diminishes responsiveness to environmental male-induction cues (from ∼90% down to ∼30%), providing a mechanistic explanation for the evolutionary decay of environmentally sensitive sex determination in NMP lineages. Nevertheless, variation in response was observed across NMP clones, suggesting the influence of secondary epistatic effects, potentially involving the other four candidate genes located within the sex-determining region (Table S2c), or reflecting the less complete suppression of *Dfh* in engineered lines relative to natural NMP clones (Figure 2e).

To further investigate the functional role of *Dfh* in male-determination pathways, we constructed a gene co-expression network based on RNA-seq data from both MP and NMP clones, sampled across key pre-adult developmental stages under the male-inducing SD photoperiod. In addition to three embryonic time points (E-12 h, E-36 h, and E-60 h), this dataset included early juvenile stages (1-day and 7-day), thereby increasing the temporal resolution and statistical power for inferring regulatory associations. Differentially expressed genes (DEGs; Table S2b) were partitioned into nine discrete co-expression modules using the Leiden clustering algorithm ^47^, with each group representing a distinct expression trajectory across developmental time.

*Dfh* was classified within Cluster 5 (Figure 2f-g), which also included eight genes involved in juvenile hormone (JH) and ecdysteroid (20E) signaling (Table S2e)—two pathways known to play central roles in *Daphnia* development and reproduction. Within this group, *Dfh* showed its strongest positive correlation with two canonical transcriptional regulators: *E75* (r = 0.84) and *HR3* (r = 0.82), both of which are downstream targets of JH and 20E signaling ^44,46^. This co-expression pattern suggests that *Dfh* may itself be a downstream effector of these endocrine pathways, a relationship that appears to be intact in MP clones but potentially disrupted in NMP clones.

To further evaluate this hypothesis, we employed AlphaFold3 ^48^ to predict protein-DNA interactions between Cluster 5 JH/20E genes and the *Dfh* upstream regions. Notably, we identified a region spanning −340 to −320 bp upstream of the *Dfh* transcription-start site that displayed strong binding affinity for both *E75* (ipTM = 0.83) and *HR3* (ipTM = 0.85) (Figure 2h). Functional validation via RNAi-mediated knockdown of *E75* and *HR3* in presumptive male embryos (from MF-treated mothers) resulted in a marked decrease in *Dfh* expression (Figure 2i; Figure S3f), providing experimental support for a regulatory hierarchy in which *Dfh* acts downstream of hormone-responsive transcription factors. Taken together, these findings suggest that *Dfh* is a key node in the developmental network governing male differentiation, likely integrating hormonal cues via direct transcriptional regulation by *E75* and *HR3*.

### A dual role of Dfh in maternal molting and sex determination in Daphnia

Sequence analysis indicates that *Dfh* encodes a putative secreted protein (Figure S4a), raising the possibility that it may contribute to the maternal transmission of environmental cues involved in sex determination. To explore this, we examined *Dfh* expression—and that of its putative hormonal regulators *E75* and *HR3*—in MP and NMP mothers carrying embryos at 12, 36, and 60 hours post-ovulation (M-12 h, M-36 h, and M-60 h). At M-60 h, a developmental stage critical for initiating the reproductive cycle and epidermal regeneration, expression levels of all three genes were significantly lower in NMP mothers relative to MP mothers (Figure S4b) under the SD condition (which induces male production), suggesting a disruption in maternal signaling pathways relevant to male-reproductive timing in NMP.

To determine the tissue specificity of *Dfh* expression in mothers, we examined bulk RNA-seq across maternal organs and found that *Dfh* is primarily expressed in the carapace and antennae (Figure 3a). We then performed single-cell RNA-seq on these tissues for mothers at M-60 h under SD conditions in both MP (KAP4) and NMP (KAP106) clones (Figure 3b). From the carapace, we recovered 10,764 cells in MP mothers and 3,602 in NMP mothers, while antennae yielded 1,146 and 1,076 cells, respectively (Figure S4c-d). Cluster analysis revealed 35 transcriptionally distinct populations in the carapace and 24 in the antennae (Figure 3c; Figure S4e). Of the 14 carapace clusters associated with cuticle formation, 13 showed significant differences in their proportional cell counts (relative to the total) between genotypes (Figure 3d; Tables S3a–b). Similarly, 9 of 10 clusters linked to molting showed different proportions. However, the proportions of antenna-cell populations were largely stable (Figure S4f; Tables S3c–d) in MP and NMP. These results localize the functional divergence between MP and NMP maternal phenotypes to specific carapace cell types.

**Figure 3:**
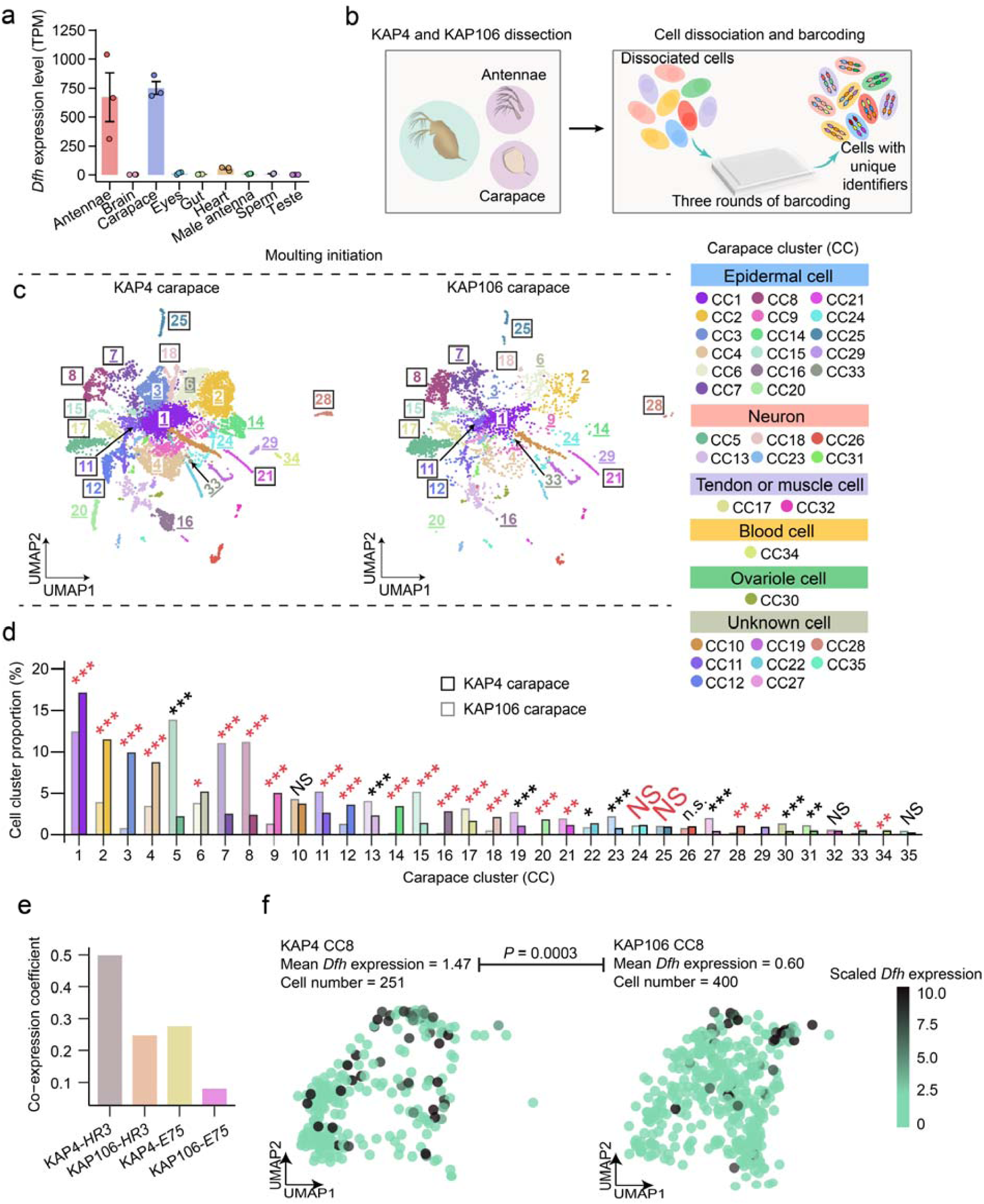
*Dfh* involvement in reproductive cycles. **a)** *Dfh* expression across different tissues. This panel shows the TPM levels of *Dfh* in nine tissues from adult KAP4 (MP) individuals. Each dot represents a biological replicate, with data presented as mean ± S.E.M. **b)** scRNA-seq workflow for carapace and antennae. Single-cell RNA sequencing (scRNA-seq) was conducted on carapace and antenna tissues from adult female KAP4 and KAP106 (NMP) individuals. Dissections were performed at a reproductive stage just before the release of current offspring, preparing for the next reproductive cycle. Dissociated cells were barcoded with unique indices before sequencing. **c)** Carapace cell clustering. Cells were clustered using the Leiden algorithm based on gene-expression profiles, identifying 35 distinct cell clusters in the carapace (represented by colored dots). Cluster annotation was based on *Drosophila* cell markers and a curated gene set involved in cuticle formation (underlined), moulting fluid regulation (black border), and other related functions. **d)** Proportional representation of carapace cell clusters normalized to total cell count of KAP4 or KAP106 carapace. A chi-square test was used to compare the proportion of each carapace cell (CC) cluster between KAP4 and KAP106 relative to the total cell count. Statistical significance is indicated as follows: NS (not significant), * (0.001 ≤ *P* < 0.05), ** (10^-6^ ≤ *P* < 0.001), *** (*P* < 10^-6^). Red font indicates clusters involvingcuticle formation or moulting fluid regulation. **e)** Co-expression of *Dfh* with *HR3* and *E75* in carapace cells. Cells with very low expression have been excluded before coefficient estimating. **f)** *Dfh* expression levels in CC8 cells. Expression levels of *Dfh* are represented by color intensity, with deeper green dots indicating higher gene expression. CC8 shows significantly higher *Dfh* expression in KAP4 compared to KAP106 (*t*-test).

Correlation analysis revealed that *Dfh* expression in MP carapace cells was more strongly positively associated with *E75* and *HR3* expression than in NMP cells, consistent with a downstream role in the hormone-responsive regulatory cascade mediating environmental sex determination (Figure 3e). One cluster in particular, CC8, accounted for 53.3% of all high-*Dfh*-expressing cells, and showed a 2.45-fold elevation in *Dfh* expression in MP relative to NMP mothers (Figure 3f). This cluster also exhibited highly expressed markers associated with epidermal remodeling and molting-fluid production (e.g., chitinase, chymotrypsinogen), implicating it in the physiological processes that preceed reproductive output (Table S3b). Comparison of differentially expressed genes (DEGs) in CC8 between MP and NMP also revealed that many of these are tied to molting-fluid production (Figure S4g; Table S3e). Intriguingly, NMP CC8 cells showed elevated expression of vitellogenin (25.75-fold) and *ovo* (2.32-fold) genes linked to yolk formation and oogenesis, implying that these same cells may be contributing to oocyte provisioning in NMP mothers.

Together, these findings support a model in which *Dfh* functions to integrate environmental signals into maternal molting and reproductive processes and has been co-opted into the embryonic sex-determination pathway. In NMP clones, reduced *Dfh* expression in specific maternal tissues may impair the relay of male-inducing environmental information to the eggs, ultimately locking development into the female pathway. This disruption could represent a key mechanistic basis for the reproductive divergence observed between MP and NMP lineages.

### Recombination Suppression Beyond Inversion

Given that the NMP-linked region harbors the sex-determination gene *Dfh* and segregates as a linked haplotype, it constitutes a sex-determining region (SDR). A pivotal event in the evolution of sex chromosomes is the onset of recombination suppression, reflected in elevated linkage disequilibrium (LD), within the SDR. Whether this process is primarily driven by sexually antagonistic selection ^19–21^ or results from a fortuitous chromosomal inversion ^18^ remains under debate.

To evaluate potential differences in LD between the *M* and *m* haplotypes in NMP clones, we performed haplotype phasing using three fully sequenced NMP genomes as reference. Phased genotypes at each locus were generated by aligning six high-quality haplotypes (from TEX58, KAP106, and NFL92) to the KAP4 reference genome. Ambiguously phased sites were excluded. This curated set of phased genotypes was then used to infer *M* and *m* haplotypes in the 67 NMP clones with high confidence. LD was quantified using the squared correlation coefficient (r²) ^49^ across different regions within the *M* and *m* haplotypes and further compared between their syntenic regions. First, in the non-SDR region (1.4–8.0 Mb) of chromosome I, LD was lowest and showed comparable levels between the *M* and *m* haplotypes (Figure 4). Second, we partitioned the SDR into the inverted region (0–0.6 Mb) and the non-inverted region (0.6–1.4 Mb). As expected, LD was markedly elevated in the inverted region in both *M* and *m* haplotypes relative to the non-SDR (Figure 4), consistent with structural suppression of recombination. Notably, LD was even higher in the *M* haplotype than in the *m* haplotype. Although NMP clones cannot intermate, they retain the ability to produce female offspring and can mate with MP clones (*mm* genotype). Consequently, the *m* haplotype can still undergo recombination at the population level. Third, in the non-inverted portion of the SDR, LD was again significantly elevated in the *M* haplotype but only slightly higher to background levels (non-SDR) in the *m* haplotype (Figure 4). Because this region lacks structural features that suppress recombination, the elevated LD in the *M* haplotype is likely maintained by additional evolutionary forces, such as selection or drift.

**Figure 4:**
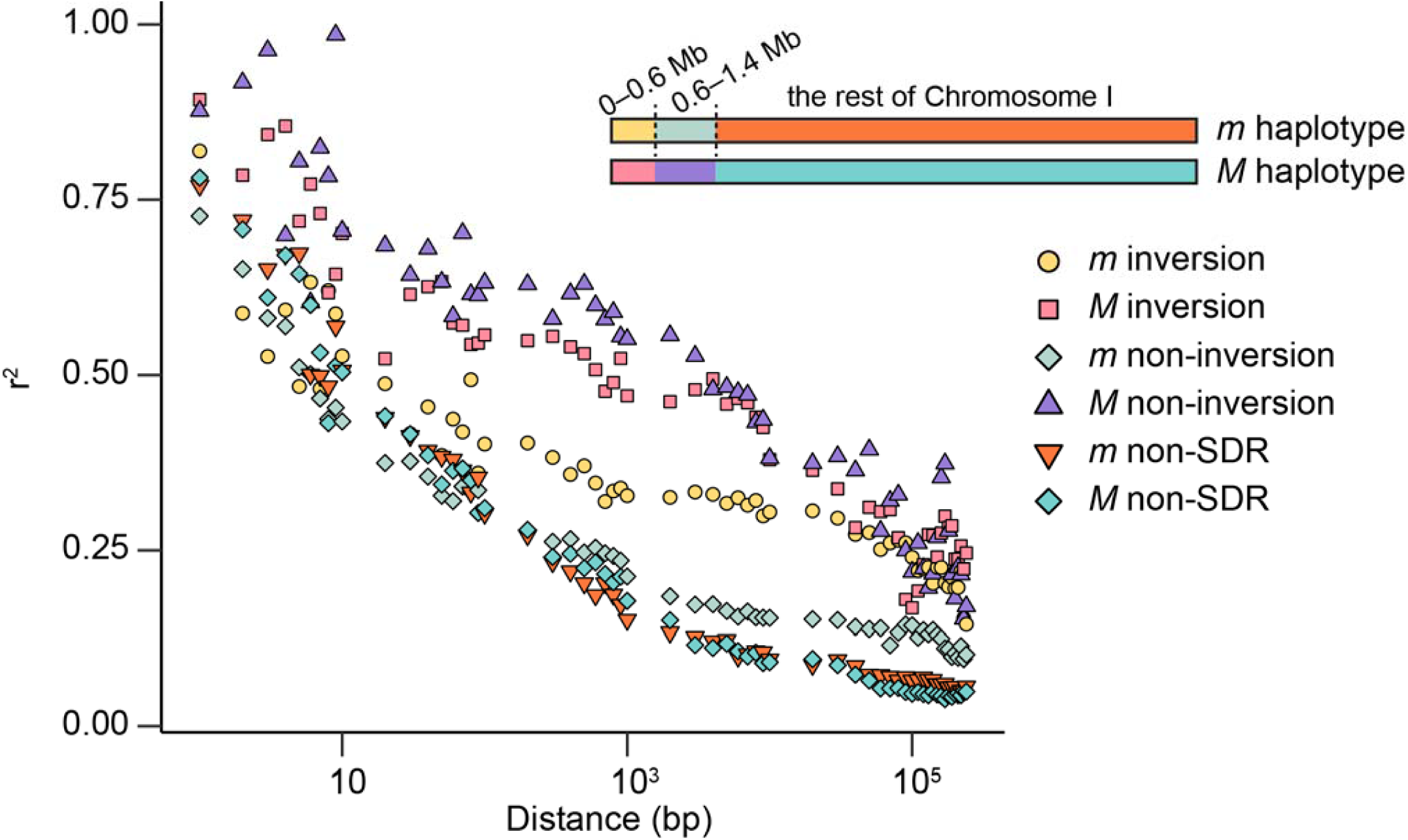
Recombination suppression beyond inversion in the SDR. We examined the decay of linkage disequilibrium (LD) across three regions: the inverted segment (∼0–0.6 Mb), the non-inverted portion of the SDR (0.6–1.4 Mb), and the flanking non-SDR region (all coordinates based on the KAP4 reference genome). Using phased genotypes from 67 NMP clones, we estimated population-level LD separately for *m* and *M* haplotypes, measured as the squared correlation coefficient (r²).

These findings indicate that the enhancement of recombination suppression within the *M* haplotype of the SDR involved a series of interacting events. Initially, inversion enhanced recombination suppression within the SDR. However, the elevated LD observed in the SDR is not solely attributable to structural constraints. Instead, other forces such as selection or genetic drift, likely act to maintain the extended haplotype structure. Given that the effective population sizes (*Ne*) of the *M* and *m* haplotypes are similar (1,582,000 vs. 1,178,000 calculated based on average πS from non-SDR region), the effects of genetic drift are expected to be comparable, making selection the more probable driver of the observed differences. This pattern aligns with classical models of sex chromosome evolution, wherein advantageous alleles, such as those promoting the NMP phenotype, become associated with non-recombining regions and are retained through selection against recombination. The high LD in NMP clones thus reflects a combined influence of structural and selective forces.

### Evolutionary origins of the SDR in NMP clones

To investigate the evolutionary origin of the SDR in *D. pulex*, we focused on two key questions: 1) how the *Dfh* gene originated and acquired its role in sex determination; and 2) how the surrounding SDR region underwent structural and functional divergence in MP clones. These two processes—the recruitment of a gene into a sex-determining role and the subsequent differentiation of the broader genomic region—represent distinct but interconnected steps in the evolution of a sex-linked genomic architecture.

The evolutionary origin of the *Dfh* locus in *Daphnia* likely occurred in the common ancestor of the *Daphnia* subgenus—after the divergence from the *Ctenodaphnia* lineage—through a series of consecutive gene duplications. This conclusion is supported by the presence of *Dfh* in multiple *Daphnia* species and its absence in *Ctenodaphnia* members such as *D. magna*, *D. lumholtzi*, *D. sinensis*, and the outgroup *Simocephalus vetulus* (Figure 5a). These findings indicate that *Dfh* emerged after the *Ctenodaphnia–Daphnia* split, which occurred at least 145 MYA ^50^. Although all *Dfh* paralogs are located on chromosome I, their distinct genomic positions suggest a history of chromosomal rearrangements (Figure 5a). Among the paralogs, only one copy has been retained in *D. mitsukuri* and members of the *D. pulex* complex. Strikingly, the copy retained in MP *D. pulex* exhibits a notably high rate of protein evolution, with a dN/dS ratio (relative to the MP outgroup *D. obtusa*) higher than 94.1% of all genes in the genome. (Figure 5b). Based on screening ∼ 3,000 clones (Table S1d), *Dfh* also exhibited relatively high πN/πS ratios among all KAP4 (MP) genes in *Daphnia* populations (Figure S5a), especially for the members of the *D. pulex* complex, including *D. arenata*, North American *D. pulex*, European *D. pulex*, *D. pulicaria*, and *D. melanica*. Only some of North American *D. pulex* clones exhibited significant NMP markers and were included in this analysis.

**Figure 5:**
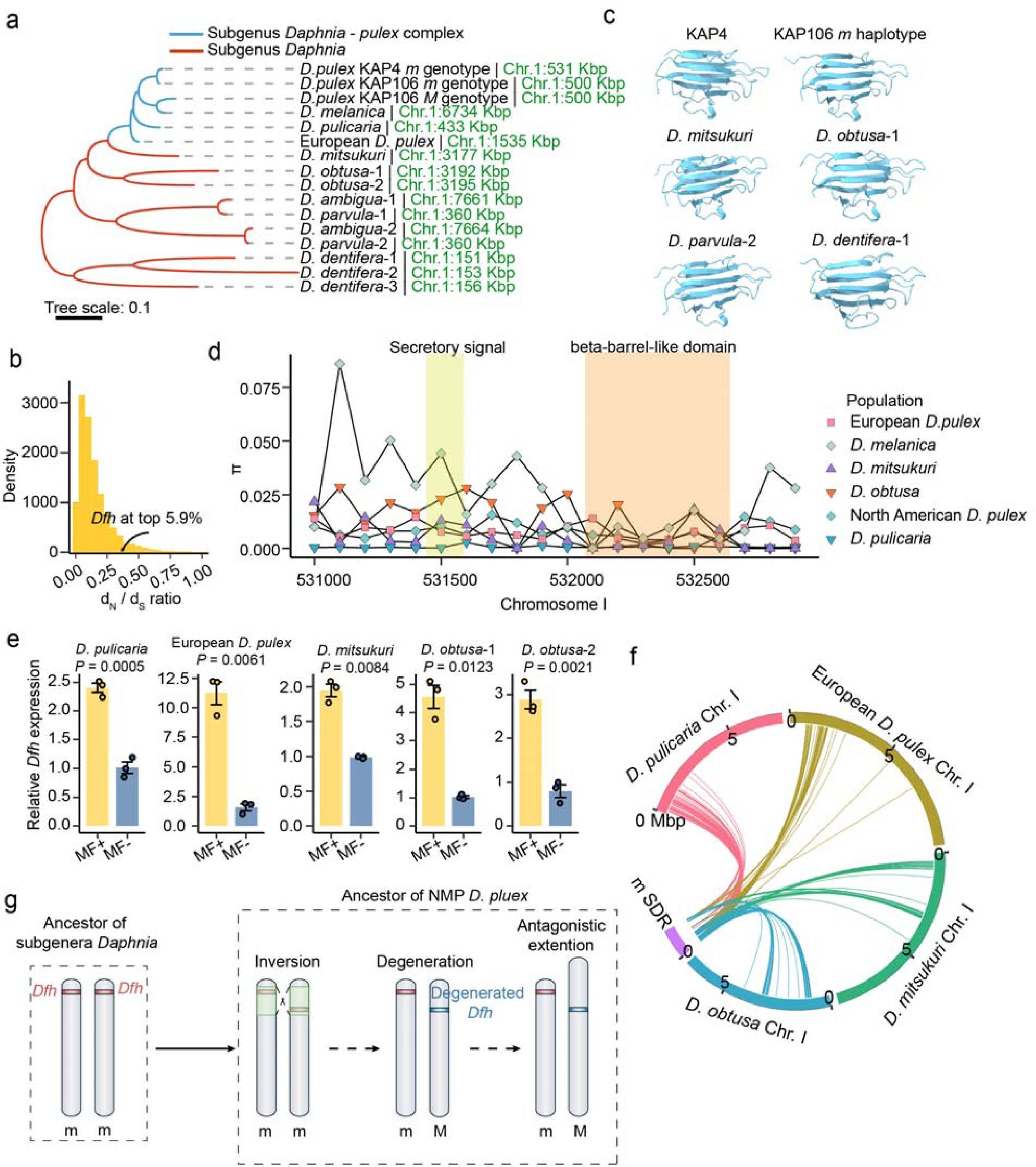
Origin and evolution of *Dfh*. **a)** Phylogenetic tree of *Dfh*. This phylogenetic tree was constructed using the Neighbor-Joining (NJ) method with *Dfh* protein sequences. The tree is consistent with species relationships. The location of *Dfh* in different species marked in green font. **b)** Distribution of d_N_/d_S_ ratios. This panel shows the distribution of d_N_/d_S_ ratios (the ratio of non-synonymous to synonymous substitutions) for all genes in *D. pulex*. The *Dfh* gene exhibits a relatively higher d_N_/d_S_ ratio. **c)** Predicted *Dfh* domain structure. Using AlphaFold3, a conserved beta-barrel-like structure was predicted across all *Dfh* homologs. This panel shows six representative examples of the conserved structure, highlighting species-wide conservation. **d)** Nucleotide diversity at the *Dfh* locus and surrounding genomic regions was estimated using 100 bp sliding windows. For each species, diversity values were averaged across populations. **e)** qPCR validation of *Dfh* expression in other *Daphnia* species. Single clones of *D. pulicaria*, European *D. pulex*, *D. mitsukuri*, and *D. obtusa* were exposed to MF for at least 2 weeks. The 60-hour embryos from over 10 individuals per species were collected for RNA extraction, and qPCR analysis was performed. In all species, MF significantly increased *Dfh* expression. Each dot represents a qPCR replicate, with data presented as mean ± S.E.M. *P*-values were obtained using t-tests comparing MF-treated (MF+) and untreated groups (MF-). **f)** Syntenic relationships between the SDR of MP *D. pulex* and Chromosome I of closely related species, with links indicating homologous gene pairs. **g)** Evolutionary model of the *M* and *m* Chromosomes. The sex-determining gene *Dfh* is thought to have originated from the ancestor of the *Daphnia* subgenus. A large chromosomal inversion in the ancestor of NMP clones suppressed recombination, leading to the accumulation of mutations and the degeneration of gene dosage on the *M* chromosome. Over time, the *M* and *m* chromosomes began to differentiate morphologically.

Despite increasing sequence divergence among *Dfh* homologs with evolutionary distance, the encoded proteins consistently maintain a conserved beta-barrel-like domain at the C-terminal (Figure 5c; Figure S5b) and a conserved secretory signal at the N-terminal (Figure S5b-c). Moreover, the beta-barrel-like domain showed lower nucleotide diversity across species than other sites (Figure 5d). These data suggest functional conservation of *Dfh* within the *Daphnia* subgenus.

To determine whether *Dfh* homologs in other species are similarly regulated by hormonal pathways, we exposed MP clones of *D. pulicaria*, European *D. pulex*, *D. mitsukuri*, and *D. obtusa* to methyl farnesoate. In all cases, *Dfh* expression was upregulated and male offspring production increased significantly (Figure 5e; Figure S6a). Across 372 clones from four *D. arenata* populations, we discovered a loss of the beta-barrel–like domain in both alleles of *Dfh* (Figure S6b). Expression assays from three representative clones revealed drastically reduced transcript levels relative to α-tubulin (Figure S6c), accompanied by a markedly low male-production rate (as low as to 17.8%; Figure S6d). For example, 6 of 14 OA15 individuals failed to produce any males over the testing period. These results suggest that *Dfh* retains a conserved hormonal-response function across the *Daphnia* subgenus, potentially mediated by its C-terminal domain (beta-barrel–like), and plays a central role in male determination. The evidence of conservation implies that hormonal control of *Dfh* is older than the inversion event, highlighting its deep evolutionary entrenchment in sex-determination pathways.

We next examined the evolutionary origin of the genetically linked SDR by analyzing genic synteny across species within the *Daphnia* genus. In species outside the *pulex* complex, such as *D. obtusa* and *D. mitsukuri*, orthologs of SDR genes are dispersed across multiple regions of chromosome I (Figure 5f). In contrast, members of the *pulex* complex exhibit these orthologs as a contiguous block, suggesting that the intact SDR structure arose in their common ancestor. Our analyses revealed recombination suppression in the SDR of both the *M* and *m* haplotypes (Figure 4), including regions unaffected by genomic inversions. To explore the basis of this suppression in *D. pulex*, we examined the genetic map constructed by Cécile *et al.* ^51^, which shows that recombination within the SDR is strongly reduced, resembling the pattern typically observed around centromeres (Figure S6f). A similar pattern was observed in its sister species *D. pulicaria* ^52^ (Figure S6f), suggesting that reduced recombination—and the emergence of the ancestral sex-linked region—arose within the *pulex* complex prior to the divergence of *D. pulex* and *D. pulicaria*. Finally, we extended our analysis to *D. obtusa*. Because no genetic map is available for this species, we used population genomic data to calculate LD (r²). The results showed similar LD decay in SDR and non-SDR regions, providing no evidence for recombination suppression in SDR in *D. obtusa* (Figure S6f).

To estimate when the inversion arose, we applied a molecular-clock approach using the equation *T = ΔS / 2μ*, where *ΔS* is the synonymous divergence between the inverted and non-inverted regions, using *D. obtusa* as an outgroup. We compared genes from the inverted and non-inverted regions to their orthologs in *D. obtusa* and calculated the average the synonymous divergence values; *ΔS* was then defined as the synonymous difference between the two regions. This analysis yielded an estimated divergence time of ∼0.19 MYA, closely matching the estimated origin of the NMP haplotype (0.19–0.85 MYA ^34^). Together, these findings suggest that the inversion arose around the same time as the earliest emergence of the NMP haplotype, marking a pivotal step in the structural evolution of the SDR.

In summary, our results suggest that *Dfh* acquired hormonally regulated functions prior to the origin of the SDR. The gene was retained as a single copy in the common ancestor of *D. mitsukuri* and the *pulex* complex and subsequently became genetically linked to the SDR during the diversification of the *pulex* complex. In the NMP lineage, a large chromosomal inversion arose, capturing *Dfh* together with additional candidate genes and neighboring loci. Regulatory and/or coding changes likely enabled *Dfh* to function as a partial genetic determinant of the sex ratio in mixed populations of MP and NMP clones. Subsequently, evolutionary forces maintained LD in alleles beyond the inversion, ultimately giving rise to the NMP phenotype and initiating the early stages of sex chromosome evolution in *D. pulex* (Figure 5g). Importantly, disruption of *Dfh* alone is insufficient to abolish male production, indicating that additional genes— such as SDR-associated DEGs (Table S2c)—also contribute to this trait and are likely maintained by selection.

## Discussion

A central question in sex-chromosome evolution is whether recombination suppression is a passive byproduct of structural changes like inversions, or if it is actively reinforced by natural selection. Our findings in *D. pulex* suggest a synergistic interplay between these forces. The SDR is located in a genomic area with ancestrally low recombination, likely due to its proximity to the centromere, a feature also observed in its sister species *D. pulicaria*. This pre-existing condition of reduced crossing over may have created a permissive environment for the evolution of a linked block of co-adapted alleles. Subsequently, a large chromosomal inversion occurred within this region, further suppressing recombination and physically locking together the genes on the nascent *M* haplotype. While these structural features provided the initial barrier, they cannot fully explain our observations. We found that LD in the *M* haplotype is significantly elevated compared to the *m* haplotype, not only within the inversion but also in adjacent non-inverted portions of the SDR. This pattern suggests that, once the NMP phenotype emerged (preventing interbreeding among NMP genotypes), natural selection acted to preserve the integrity of the entire advantageous *M* haplotype by purging recombinant haplotypes that would otherwise disrupt favorable allele combinations. Although genetic drift could theoretically contribute, the similar effective population sizes of the *M* and *m* haplotypes make it unlikely to play a major role. Moreover, additional candidate genes associated with the NMP trait were identified in the non-inverted portions of the SDR; these genes may be maintained in linkage with *Dfh* by natural selection, jointly contributing to the trait’s function. This finding aligns with theoretical predictions that LD could be maintained by selection that favors reduced recombination between sex-determining loci and their surrounding alleles ^23,53^. Empirical studies in animals also support this interplay between selection and structure. For instance, in *Poecilia reticulata* (guppies), populations under stronger sexual selection display more extensive X–Y recombination suppression without corresponding large inversions ^54^. In *Mus musculus domesticus*, suppressed recombination around sex-determining loci extends beyond inversion boundaries, suggesting selective maintenance ^55^. Songbirds and sticklebacks further demonstrate that accumulation of sexually antagonistic genes around sex loci often assembles prior to or concurrent with structural rearrangements ^56,57^. Consistent with this, we identified several additional DEGs between NMP and MP within the SDR that may contribute to sex-specific functions and warrant further investigation.

More broadly, these findings fit within the framework synthesized by Zhou *et al.*, who highlight how dosage compensation, epigenetic regulation, and 3D chromatin architecture evolve under sexual conflict and selective pressures ^6^. Lenormand & Roze specifically propose a “regulatory model,” where selection on gene regulation is actively shaped by recombination suppression, often caused by structural changes ^24^. Our results from *Daphnia* provide a rare empirical case that directly links structural changes with selection during the earliest stages of sex-chromosome evolution. While the inversion likely established the initial recombination barrier, natural selection appears to have expanded and reinforced suppression into neighboring regions—a synergy that underpins proto–sex chromosome evolution. Together, these results underscore selection’s pivotal role in establishing early sex-chromosome differentiation.

Another key question in the origin of sex chromosomes is when and how sex-determining genes (SDGs) emerge—whether they arise after structural changes such as inversions that facilitate their recruitment, or if their sex-determining functions predate these structural rearrangements. Our study in *D. pulex* provides empirical support for the latter scenario: SDGs may acquire functional roles prior to the onset of recombination suppression and structural divergence. First, the *Dfh* gene originated within the *Daphnia* subgenus and acquired a conserved function in hormonally regulated pathways, as evidenced by its upregulation by methyl farnesoate across multiple species. This ancestral role in modulating male developmental processes predated the evolution of the NMP trait. Second, in the common ancestor of the *pulex* complex, genes that now constitute the SDR became assembled into a contiguous genomic block. Finally, approximately 0.19 MYA, a large chromosomal inversion occurred, capturing a specific allele of *Dfh* (*Dfh-M*) and neighboring loci. This event coincided with the origin of the NMP haplotype and likely stabilized the genetic basis for a transition from purely environmental to partially genetic sex determination. Comparable evolutionary patterns have been reported in other taxa. In medaka (*Oryzias latipes*), the SDG *dmrt1bY* arose by duplication and acquired sex-determining functions before recombination suppression evolved on the Y chromosome ^58^. In platyfish (*Xiphophorus maculatus*), recombination suppression expanded gradually around preexisting sex loci via the accumulation of sexually antagonistic variants and local rearrangements ^59^. Similarly, Zhou *et al.* showed that in multiple vertebrate lineages, ancient transcription factors were co-opted for sex determination before large-scale chromosomal changes occurred ^6^. Together, these findings—including our results in *Daphnia*—support a general model in which functionally competent SDGs can emerge and gain partial or full regulatory roles within recombining autosomes, and are only later embedded within structurally and functionally differentiated proto-sex chromosomes. In this model, recombination suppression such as inversions are secondary events that help stabilize or differ the SDG’s effects by preventing recombination, thereby catalyzing the early stages of sex chromosome evolution.

Our results demonstrate that *Dfh* plays a central role in sex determination in *D. pulex*, as CRISPR-induced indels dramatically reduced male production rates, underscoring its essential contribution to male induction. Consistently, evidence from *D. arenata* populations highlights the importance of *Dfh*’s β-barrel–like domain: loss of this domain was associated with a drastic decline in male production (Figure S6c-d), with some clones producing no males at all—for example, in population OA15, 6 of 14 clones failed to produce males within the testing period (Figure S6d). These findings confirm that the structural integrity of *Dfh* is indispensable for its function and that disruption of this gene severely compromises environmentally induced male production. However, the reduction in male production from ∼90% to ∼30% in CRISPR-induced KO lines and to ∼20% in β-barrel–like domain–deleted *D. arenata* populations suggests that secondary genes within the SDR act alongside *Dfh*, accounting for the remaining 20–30% of sex determination; combined loss of these factors could abolish environmental sex determination entirely. Transcriptomic analyses identified four additional candidate genes (Table S2c) within the SDR that are differentially expressed between MP and NMP clones under male-inducing conditions. Notably, two candidates are located within the inversion, whereas the other two reside in the non-inverted portion of the SDR, potentially associated with the selection on the linkage maintain between the inversion and the non-inverted region. Two of four genes are completely characterized, one is *leucine-rich PPR motif-containing protein, mitochondrial-like*, and the other is *Kinase suppressor of Ras 1* (*ksr1*; LOC124192329). *ksr1*, positions in the non-inverted region and includes 9 NMP markers in *M* allele, is particularly noteworthy. The gene *ksr1* functions as a scaffold protein that facilitates Raf/MEK/ERK complex formation and Ras/MAPK signaling ^60,61^, a pathway broadly implicated in cell fate decisions and sex-determination cascades ^62^. This raises the possibility that *Dfh* interacts epistatically with SDR-linked genes such as *ksr1*, with natural selection maintaining their linkage. Analogous to the *Drosophila* Sex-lethal (*Sxl*) pathway, in which *Sxl* recruits *Unr* (an upstream regulator of Ras) to regulate sex-specific mRNA translation ^63,64^, sex determination in *Daphnia* may likewise depend on coordinated networks of regulatory and signaling genes. Collectively, these findings suggest that the NMP phenotype has a polygenic basis, with *Dfh* as the principal determinant but additional SDR-associated genes contributing to the stability of the trait.

Therefore, it is highly probable that the NMP phenotype arises from the cumulative or epistatic effects of reduced *Dfh* expression in concert with regulatory or functional changes in these other linked genes. This polygenic architecture provides a compelling rationale for the strong selective pressure to maintain the *M* haplotype as an intact unit. Suppressed recombination would be highly advantageous, as it prevents the co-adapted suite of alleles responsible for the NMP trait from being dismantled by crossing over.

The types of genes that are predisposed to evolve into SDGs remains an open question. While classical models emphasize mutations in existing components of conserved sex-determination cascades, accumulating evidence suggests that SDGs can also emerge from novel genes or through regulatory innovation. In *D. pulex*, *Dfh* appears to have been co-opted into the hormonal regulatory network downstream of JH and 20E signaling ^44,46^, yet its regulation seems to be independent of the canonical arthropod sex-determination gene *doublesex*, which is also the downstream of hormonal regulation ^6^. This decoupling highlights that SDGs can be integrated into sex-determination pathways through regulatory rewiring, without originating from the core cascade itself. Interestingly, a gynodioecy-like system has also evolved independently in *D. magna*, but it is located on a different chromosome and involves a distinct set of genes, suggesting the emergence of a distinct SDG in *D. magna* ^50,65^. These cases illustrate that proto-sex chromosomes can evolve independently in different lineages under similar selective pressures. Together, these findings support a flexible model of SDG origin, where regulatory potential, ecological context, and structural changes like inversions interact to facilitate the emergence of genetic sex-determining systems. Although the specific genes involved may vary, the broader evolutionary trajectory often converges directly or indirectly on conserved downstream elements such as *doublesex*, reflecting a common endpoint despite diverse origins.

What mechanisms allow clones carrying a proto-sex chromosome to persist within a population? The stable maintenance of proto-sex chromosomes, particularly in systems like *D. pulex* where a gynodioecy-like mating system is present, involves a complex interplay of genetic, ecological, and evolutionary factors. First, *D. pulex* have been shown to exhibit inbreeding depression ^66–69^. The coexistence of genetically distinct NMP and MP clones may facilitate outcrossing, since NMP clones lack the ability to self-fertilize. This promotes genetic exchange and reduces the risk of inbreeding depression, thereby preserving genetic diversity in the population. Second, negative frequency-dependent selection may stabilize the polymorphism. Once NMP clones invaded a population in which MP clones are common and males are abundant, they avoid the costs of male production, benefiting from increased reproductive efficiency. Conversely, when NMP clones dominate, MP clones gain a relative advantage by providing the necessary males for sexual reproduction. This dynamic can prevent either clone type from fixing in the population. Together, these findings suggest that the maintenance of clones carrying proto-sex chromosomes does not solely depend on intrinsic genetic factors like the inversion or SDG, but also on broader ecological and evolutionary contexts that stabilize such systems via selection and mating system dynamics.

## Data availability

The data generated in this study have been deposited in the NCBI’s short read archive database under PRJNA1003069 (RNA-seq), PRJNA1314279 (scRNA-seq) and PRJNA988494 (population genomes).

## Acknowledgements

We thank Emily Williams, Tarah Schaffer, Andrew Zhao, Andres Rios Munoz, Om Gawali, Spencer Collins, George Bcharah, Aram Nejad, and Jared Alvarez for their assistance with *Daphnia* maintenance and experiments. We are also grateful to Michael E. Pfrender, Ken Spitze, Chris Steiner, Adam Petrusek, Xiaolin Ma and Brooks Miner for providing *Daphnia* clones. Special thanks to Andrew C. Zelhof for his assistance with microinjection. This work was supported by grant from the National Institutes of Health (R35GM122566), the National Science Foundation (DBI-2119963 and IOS-1922914) to Michael Lynch, and by the National Natural Science Foundation of China (Grant 32471695) to Zhiqiang Ye.

## Author contributions

W.W. and Z.Y. designed research; W.W., M.L., T.B., R.S., Z.Y., Y.H., S.N., L.W., S.X., H.L. and E.K. performed experiments; L.G. and B.L.C. collected and maintained *Daphnia*; W.W. and Z.Y. conducted bioinformatic analyses; W.W. and Z.Y. analyzed data; and W.W., Z.Y. and M. L. wrote the paper.

## STAR★Methods

### Key resources table

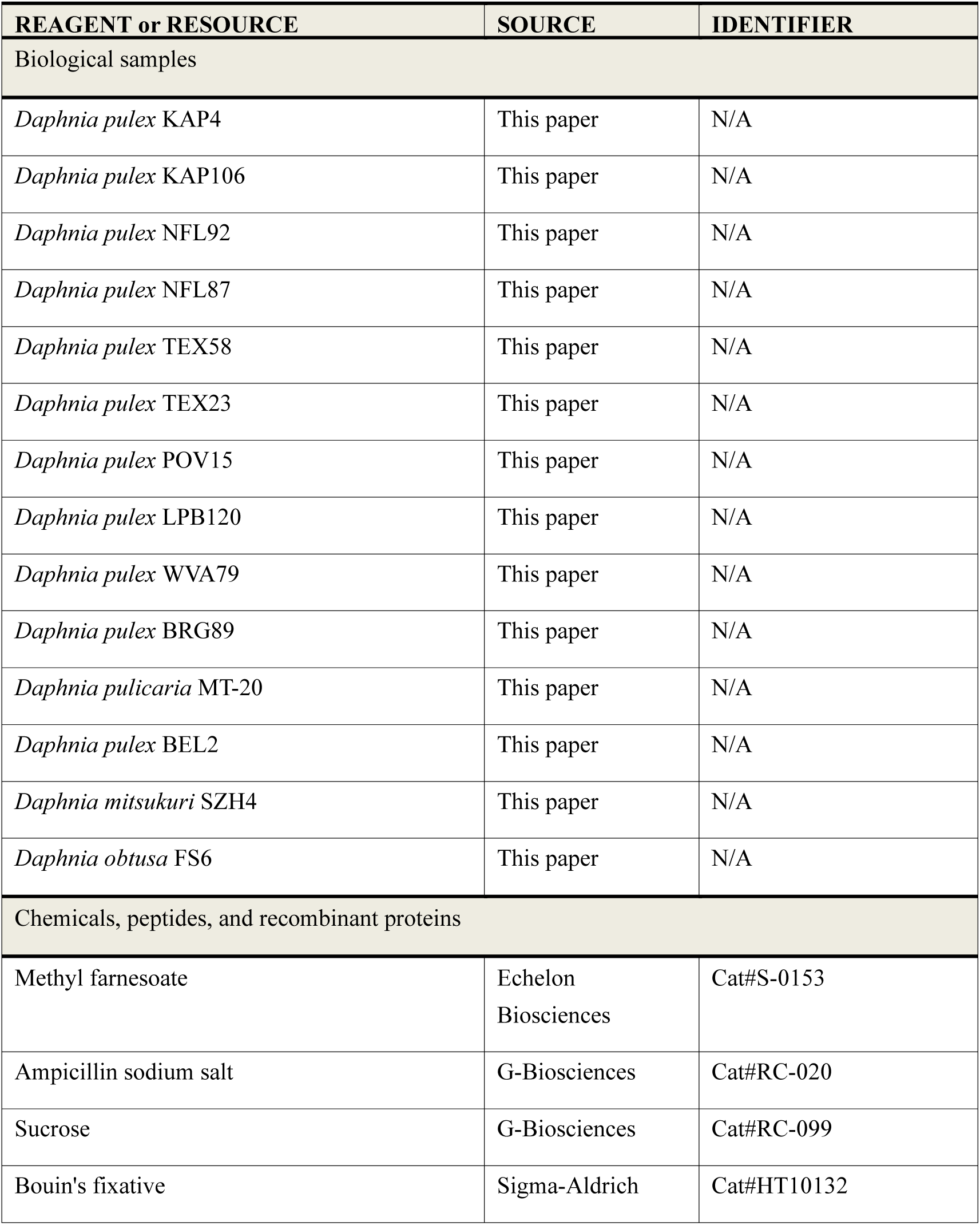
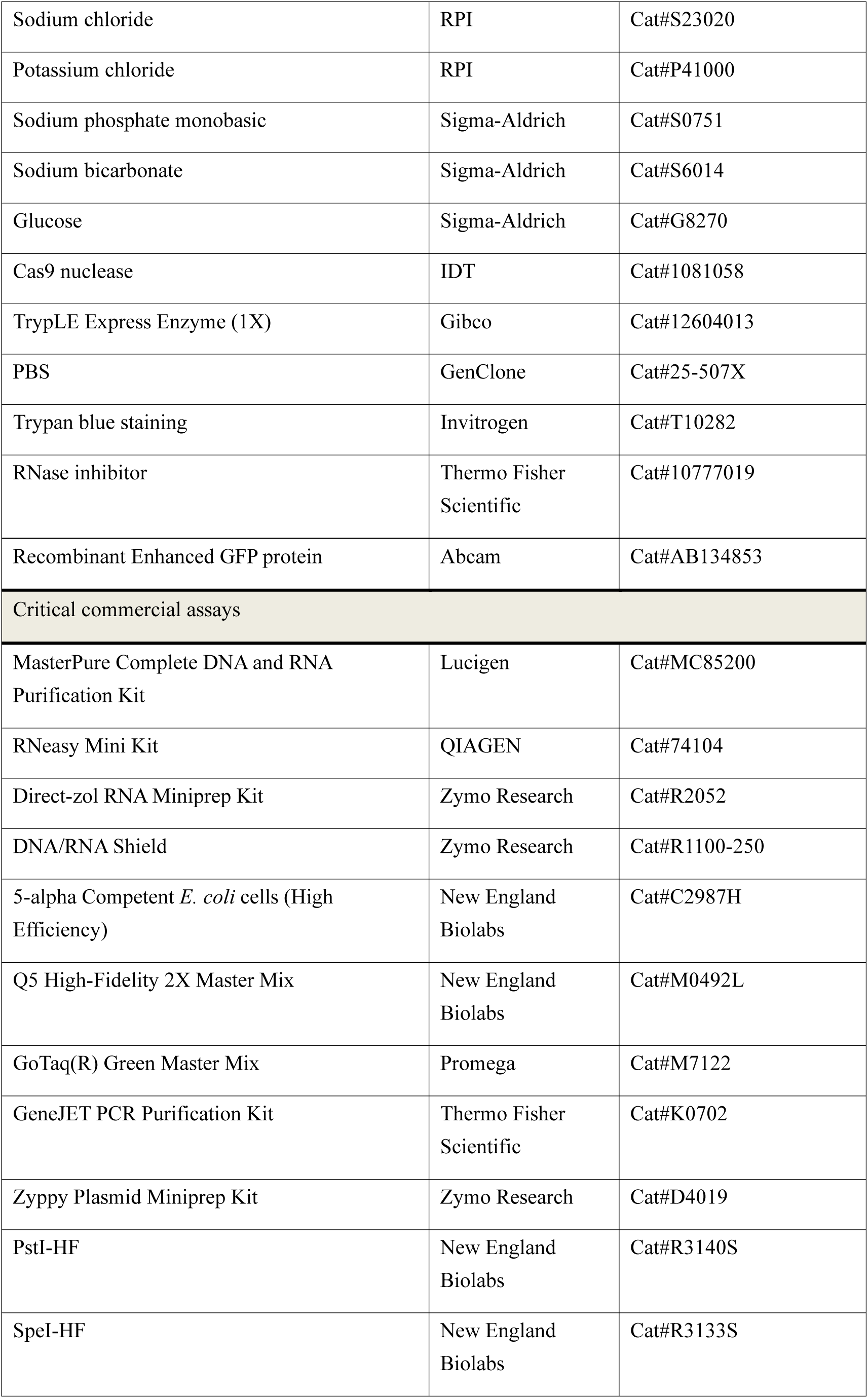
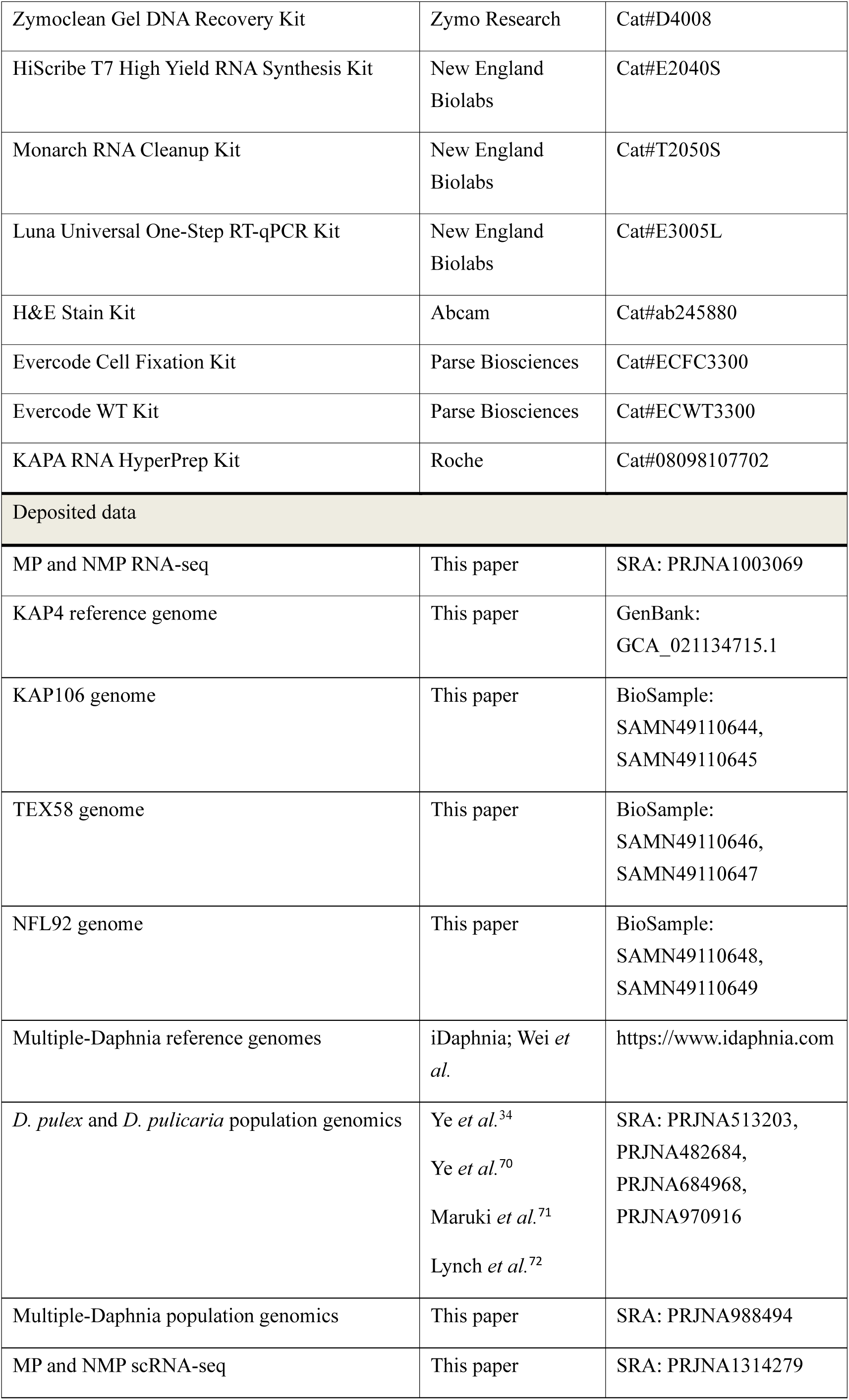
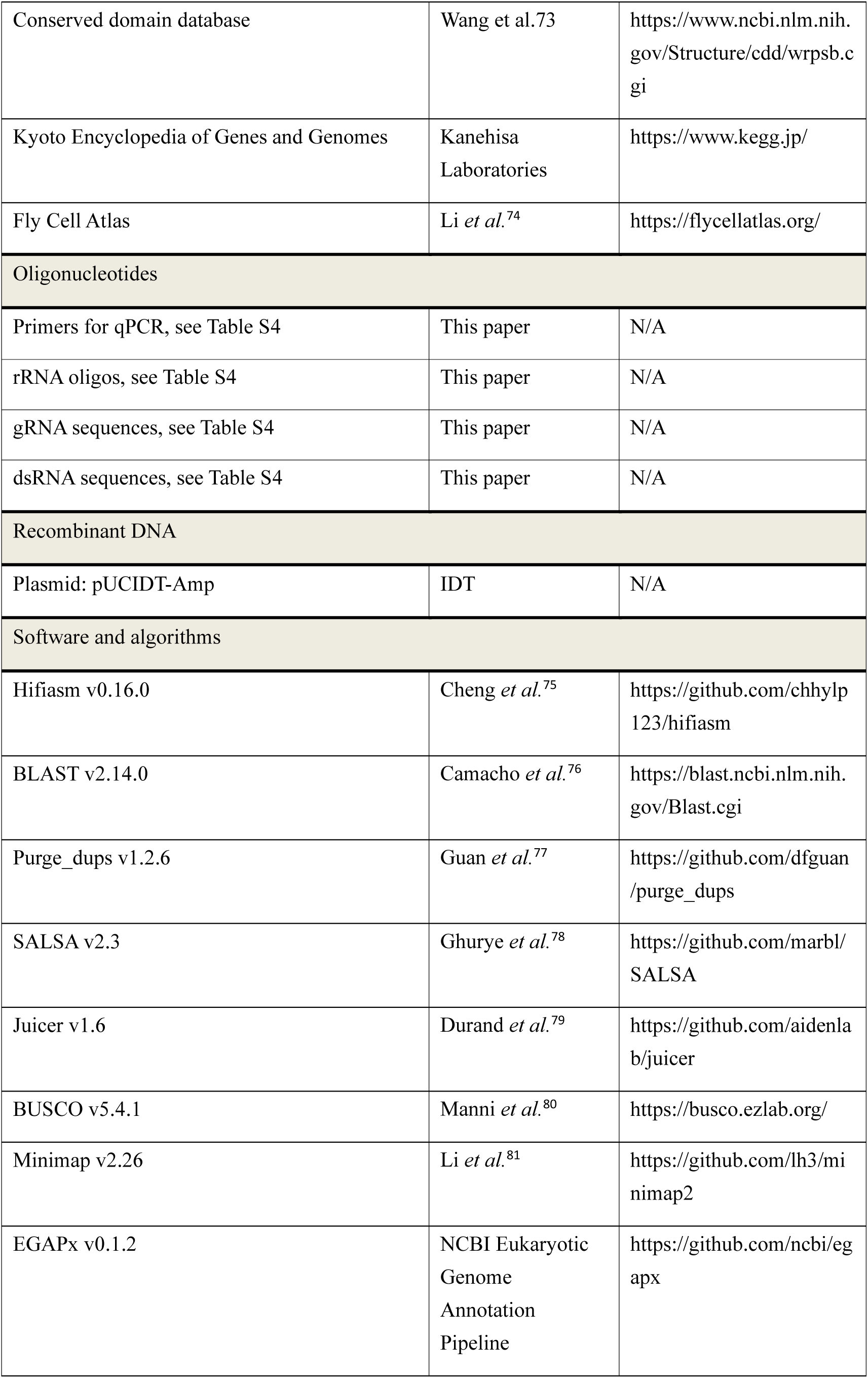
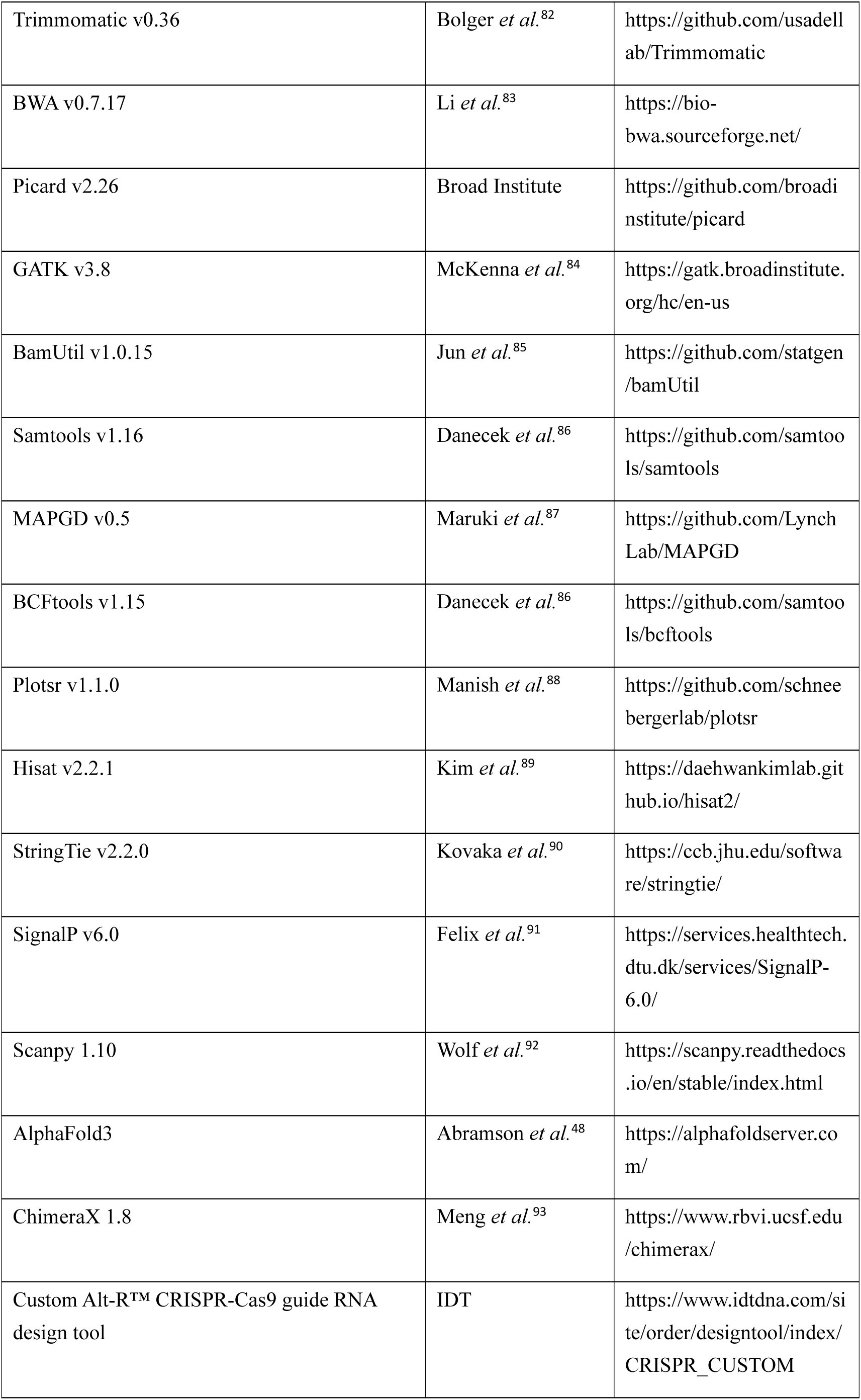
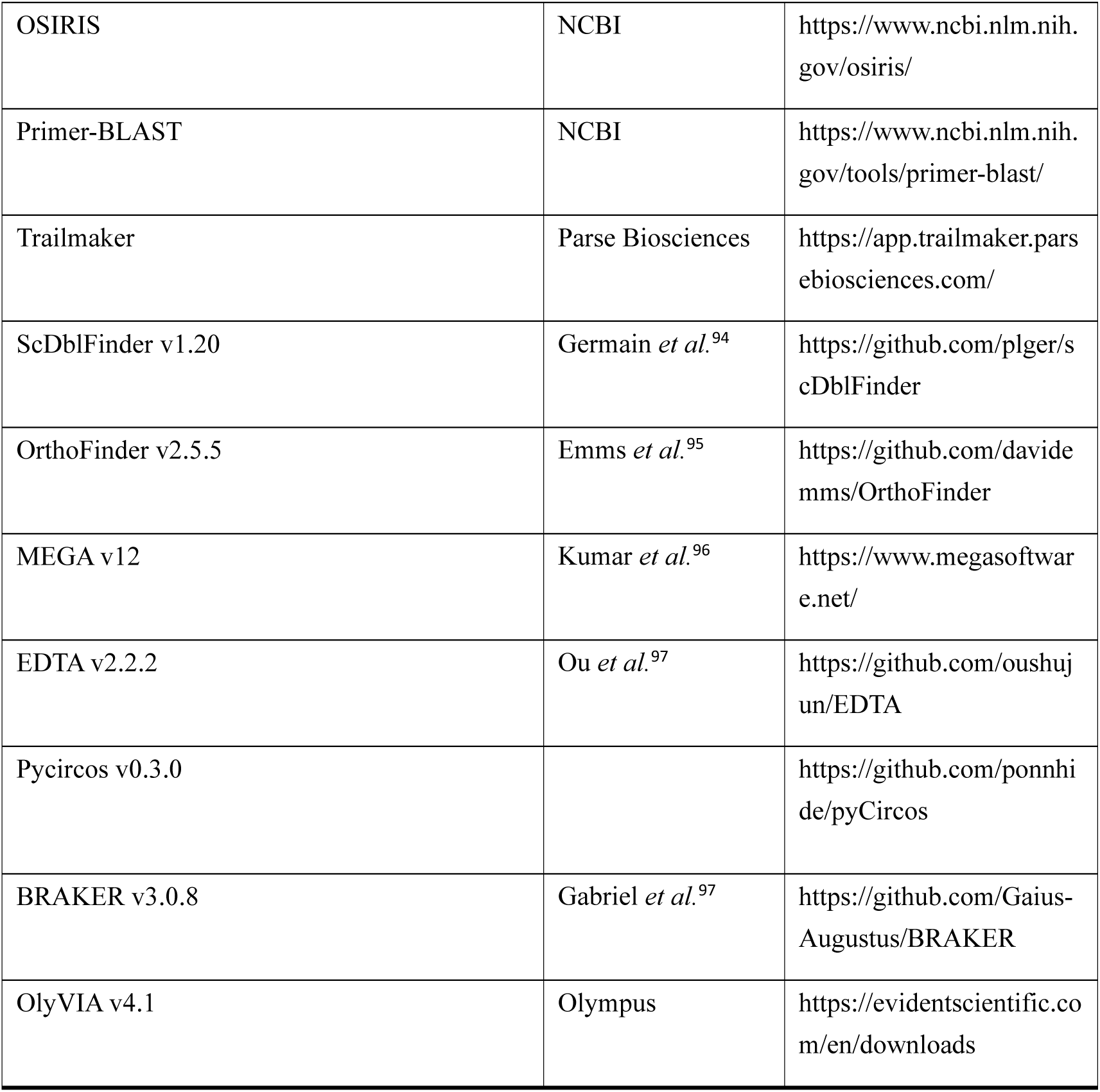

## Method details

### Sample collection

*Daphnia* samples were collected from the United States between 2013 and 2023 (Table S1d). To ensure that each isolate likely originated from a distinct resting egg, we collected early adult individuals from the water column prior to the onset of reproduction. These isolates were then clonally propagated and maintained in the lab.

### Generating genome assemblies for MP and NMP clones

A total of 167 clones from *D. pulex* populations (KAP, POV, TEX, LPB, and NFL) were tested for their male-producing ability by inducing male production through methyl farnesoate (MF) treatment ^34^. From these, we selected one male-producing (MP) clone, and three non-male-producing (NMP) clones, KAP106, NFL92 and TEX58 for genomic sequencing. To get genomic DNA, at least 500 adults from each clone, derived from a single female, were collected. The individuals were thoroughly surface-sterilized by rinsing with autoclaved lake water. Genomic DNA was extracted using the MasterPure Complete DNA and RNA Purification Kit (Lucigen), with DNA quality and integrity assessed using a NanoDrop spectrophotometer (Thermo Fisher Scientific), Qubit 4 Fluorometer (Invitrogen), and 4200 TapeStation (Agilent Technologies). DNA was collected in DNA LoBind tubes (Eppendorf) to ensure maximal nucleic acid recovery.

High-quality DNA from KAP4 and KAP106 clones was submitted to UC Berkeley QB3 Genomics (California, the United States) and UC Davis Genome Center (California, the U.S.), respectively, and sequenced using the PacBio Sequel II platform. This generated 13.72 Gb and 14.00 Gb of high-precision HiFi reads, with average read lengths of 10.87 kb and 16.30 kb for KAP4 and KAP106, respectively. Hi-C data for KAP4 and KAP106 were generated from over 50 adults. The Hi-C library construction was performed by Phase Genomics (Washington, the U.S.), and the resulting libraries were paired-end sequenced using the Illumina platform (∼200× depth coverage). High-quality DNA from the NFL92 and TEX58 clones was sent to the Arizona Genomics Institute (Arizona, the U.S.) for sequencing using the PacBio Revio platform. This produced 33.22 Gb and 27.18 Gb of high-precision HiFi reads, with average read lengths of 12.68 kb for NFL92 and 12.36 kb for TEX58.

The primary KAP4 genome was assembled using the default parameters of Hifiasm v0.16.0 ^75^. To ensure accuracy, we removed contaminated contigs from bacteria and algae by comparing the primary genomes to 2,000 randomly selected bacterial genomes and 130 algae genomes, which were downloaded from NCBI, using BLAST v2.14.0 ^76^ (identity > 95%, alignment length > 500, and E-value < 10^-5^). Haplotigs and contig overlaps were eliminated with Purge_dups v1.2.6 ^77^, and scaffolding of the cleaned contigs was completed by Hi-C data using SALSA v2.3 ^78^. To further link scaffolds, Hi-C maps were generated using Juicer v1.6 ^79^, followed by manual correction of assembly errors. Additionally, genetic maps of *D. pulex* obtained from Molinier *et al* ^51^. were used to anchor genetic markers to the KAP4 genome, validating the manually corrected assemblies. This process ultimately resolved 12 chromosomes for the KAP4 (133.2Mb) reference genome, with a BUSCO v5.4.1^80^ completeness score of 98.2%.

The primary haplotype-resolved KAP106 genome was assembled using Hifiasm with default parameters, integrating PacBio and Hi-C data. NFL92 and TEX58 were assembled solely from PacBio data. Haplotigs and contig overlaps were removed using Purge_dups, yielding near chromosome-level diploid genomes, typically with 1–2 large scaffolds per chromosome. To position these scaffolds, we aligned the NMP assemblies to KAP4 using Minimap2-2.26 ^81^. This process ultimately resolved 12 chromosomes for the NMP haplotypes, with BUSCO completeness scores from 95.2% to 98.4%.

### Performing functional and structural annotation

To gain transcriptome-based insights into gene expression, full-length transcriptome sequencing (Iso-Seq) and RNA-seq profiles were generated for KAP4 and KAP106 following exposure to various environmental challenges, including temperature shifts, changes in photoperiod, and Methyl farnesoate (MF) exposure. *Daphnia* were subjected to 1) a long-day photoperiod (14 hours light/10 hours dark) at 10°C, 18°C, and 24°C; 2) a short-day photoperiod (10 hours light/14 hours dark) at 18°C; and 3) exposure to 800nM MF (Echelon Biosciences) for 48 hours under long-day conditions (18°C).

Approximately 20 adults were maintained per beaker with mixed COMBO ^98^ and lake water, fed every two days with 150,000 cells of *Scenedesmus obliquus* per mL. For each environmental condition, over 50 adults were collected and rinsed with RNase-free water. Total RNA was extracted using either the RNeasy Mini Kit (QIAGEN) or the Direct-zol RNA Miniprep Kit (Zymo Research). RNA concentrations were measured with a NanoDrop spectrophotometer, and RNA from five environmental conditions was pooled equally. The quality and integrity of the pooled RNA were assessed using the NanoDrop spectrophotometer and the 4200 TapeStation (RIN > 8).

Full-length Iso-Seq was conducted at UC Berkeley QB3 Genomics using the PacBio Sequel II platform (one flowcell per sample). *Daphnia*-specific rRNA oligos (Table S4) were synthesized by Integrated DNA Technologies (Iowa, the U. S.), and total RNA was submitted to the Tgen Collaborative Sequencing Center (Arizona, the U.S.) for rRNA depletion using the KAPA RNA HyperPrep Kit (Roche). The sequencing was performed on the Illumina NovaSeq platform.

Short-read and long-read sequencing data for KAP4 were submitted to NCBI and used to annotate the *D. pulex* reference genome using the Eukaryotic Genome Annotation Pipeline (EGAPx v0.1.2). The final KAP4 annotation is available through NCBI at: https://www.ncbi.nlm.nih.gov/genome/annotation_euk/Daphnia_pulex/100/. To annotate both haplotypes of the KAP106 genome, we used BRAKER v3.0.8 ^99^ with input from both short-read and long-read sequencing data. For the TEX58 and NFL92 haplotypes, annotations were generated using BRAKER based on short-read RNA-Seq data collected from multiple developmental stages (see below). Additionally, transposable elements (TEs) in each genome or haplotype were identified using EDTA v2.2.2 ^100^ with default parameters.

### Comparison of genome structures and genes

To identify structural variations between MP and NMP clones, NMP *M* haplotypes were aligned to the *m* haplotypes using Minimap2. The resulting BAM files were indexed with Samtools v1.16.1 ^86^, and genomic synteny among NMP haplotypes and KAP4 was analyzed using Plotsr v1.1.0 ^88^. Only large-scale structural features— syntenic blocks, inversions, translocations, and duplications—were considered, using a minimum alignment length threshold of 20 kb of matching bases.

To delineate shared gene content in the *M* haplotypes, we first searched for homologues of KAP106 *M*-haplotype genes in the TEX58 and NFL92 *M* haplotypes. Reciprocal best-BLAST hits were obtained i) between KAP106 *M*-haplotype genes and the annotated *M*-haplotype genes of TEX58 and NFL92 and ii) between KAP106 coding sequences and the raw genomic DNA of TEX58 and NFL92, thereby minimizing false negatives due to annotation errors. Genes detected in all three *M* haplotypes by either comparison were designated common *M*-haplotype genes; for multi-copy families, we retained the minimal orthologous set shared across the three haplotypes.

To identify the putative progenitor copy of novel duplications, common *M*-haplotype sequences (from KAP106) were queried against the KAP4 reference with BLAST, retaining top hits showing ≥ 70 % identity, ≥ 50 % alignment coverage, and E-values < 10⁻⁶. A gene was classified as duplicated in the *M* haplotype when no orthologue—or a lower copy number—was found in the corresponding NMP region of KAP4. Conversely, gene losses were detected by BLASTing KAP106, TEX58 and NFL92 *M*-haplotype gene sets against the KAP4 genome; any gene present in the KAP4 NMP region but absent from all three *M* haplotypes was considered lost.

### NMP and MP genotype analysis

To identify NMP-specific genotypes (NMP markers), we analyzed 67 NMP and 100 MP clones from five *D. pulex* populations (KAP, POV, NFL, TEX, and LPB; BioProject accession number: PRJNA513203). Raw sequencing data were cleaned using Trimmomatic v0.36 ^82^ to remove adapter sequences and trim low-quality reads. The cleaned reads were then mapped to the KAP4 reference genome using BWA v0.7.17 ^83^, allowing the alignment of both paired and unpaired reads. Merged and sorted alignments were processed using Picard v2.26 (https://broadinstitute.github.io/picard/) to remove PCR duplicates with default parameters. Realignment of filtered reads around indels was performed using GATK’s “RealignerTargetCreator” and “IndelRealigner.” (v3.8) ^84^. We further clipped overlapping reads using BamUtil v1.0.15 ^85^ (https://github.com/statgen/bamUtil) and generated pileup files using Samtools v1.16 ^86^.

Allele frequencies were estimated across the *Daphnia* population using MAPGD v0.5 ^87^, followed by genotype calling with stringent filtering criteria. For each site within a given clone, genotypes were classified as homozygous or heterozygous based on the sequencing depth of the four nucleotides (A, G, C, and T). Alleles were ranked by read depth as follows: the major-depth allele (highest depth), the minor-depth allele (second highest), and putative sequencing errors (third and fourth alleles, which are unlikely under diploidy). To ensure reliability, we required the total read count of the major and minor alleles to be ≥ 6, and the read count for each error allele to be ≤ 3. A homozygous genotype was assigned if the minor allele count was 0. To evaluate whether the major and minor allele counts were consistent with a heterozygous genotype, we tested for fit to a binomial distribution, applying a Bonferroni correction for multiple comparisons across testing clones. Sites with adjusted *P*-values < 0.05 were classified as homozygous. For a given site, genotypes observed in fewer than 20 MP or NMP clones—regardless of zygosity—were excluded from further analysis.

To identify consensus genotypes specific to NMP clones, we compared genotype patterns between NMP and MP groups, requiring that candidate genotypes be absent from MP clones, thereby representing potential markers associated with the NMP phenotype. Genotypes observed in three or fewer clones were considered rare and excluded. To obtain haplotype-resolved NMP markers, we aligned both haplotypes of the KAP106 genome to the KAP4 reference genome using Minimap2. Haplotype-resolved variants were then called using BCFtools v1.15 ^86^. This allowed us to phase NMP markers onto each haplotype, providing a more detailed understanding of the genetic architecture linked to the NMP phenotype.

### Other *Daphnia* populations

Individual *Daphnia* clones were isolated into separate beakers in the lab and cultured until reaching sufficient densities for DNA extraction. Genomic DNA was extracted using the MasterPure Complete DNA and RNA Purification Kit (Lucigen). Approximately 1000 clones from 16 populations—including North American *D. pulex*, European *D. pulex*, *D. arenata*, *D. melanica*, *D. pulicaria*, *D. mitsukuri*, and *D. obtusa*—were sequenced. These samples were sent to the Center for Genomics and Bioinformatics at Indiana University (Indiana, the U. S.) or TGen for paired-end sequencing (100 or 150 bp) on Illumina platforms such as NextSeq, HiSeq, or NovaSeq, generating an average of 10X coverage. Additionally, ∼1500 clones from 15 N. A. *D. pulex* populations and 3 *D. pulicaria* populations were downloaded from previous studies ^34,70–72^ (BioProject accession number: PRJNA482684, PRJNA513203, PRJNA684968, and PRJNA970916).

Raw sequencing data were cleaned using Trimmomatic to remove adapter sequences and trim low-quality reads. The cleaned reads were then mapped to the KAP4 reference genome using BWA. Genotypes were estimated to use the same pipeline in analysis of MP/NMP genotypes.

### Nucleotide diversity analysis

We define πmm and πMm as the average per-site nucleotide heterozygosities within the MP and NMP populations, respectively. For each population, we calculated nucleotide heterozygosity (π) at these neutral sites using the formula 2pq, where p represents the minor allele frequency at the site within the population. Sites where more than 70% of clones have missing genotypes (due to low coverage or sequencing errors) were excluded from the analysis.

Additionally, we calculated the nucleotide heterozygosity of *M* haplotypes (πMM). To compute the average nucleotide diversity between alleles carried by pairs of NMP (*Mm*) isolates (π*), we used a four-allele comparison. For instance, if one NMP clone has the genotype *AB* and another clone has genotype *CD*, we compare *A* to *C*, *A* to *D*, *B* to *C*, and *B* to *D*, and the heterozyosity at the site is the fraction of these four pairs that are heterozygotes. Given that:

π^∗^=0.25π_MM_+0.5π_Mm_+0.25π_mm_, where π_mm_ and π_Mm_ represent the average site-specific heterozygosity within *mm* (MP) and *Mm* (NMP) clones, respectively. For each gene, we calculated the average per-site diversity (π_mm_, π_Mm_ and π*). Then, π_MM_ of each gene can be estimated by π_MM_=4π_∗_−π_mm_−2π_Mm._

To estimate the ratio of π_N_ (nucleotide heterozygosity at 0-fold amino-acid replacement sites) to π_S_ (nucleotide heterozygosity at 4-fold redundant sites), we computed π_N_ and π_S_ for both MP and NMP populations. 4-fold redundant sites in genes—regions traditionally viewed as being under minimal selective constraint. We focused exclusively on those 4-fold redundant or 0-fold amino-acid replacement sites that did not overlap with other functional categories. To calculate π_N_/π_S_, at least 3 genotyped sites per gene are required to generate reliable estimates of π_S_.

### RNA-Seq of MP and NMP clones across developmental stages and photoperiods

To perform RNA-Seq, we selected one MP and one NMP clone from each of five *D. pulex* populations: NFL, TEX, LPB, POV, and KAP. The male-production ability of five MP clones (NFL87, TEX23, LPB120, POV15, KAP4) and three NMP clones (NFL92, TEX58, KAP106) had been validated in a previous study. Due to the lack of surviving NMP clones from the LPB and POV populations in the laboratory, we screened NMP markers of hundreds of clones from three additional *D. pulex* populations, BRG (collected in the U.S., 2014), WVA (U.S., 2014), and SPS (U.S., 2013) (Table S1d), to identify alternative NMP candidates. Then we tested some clones from WVA and BRG, which show significant NMP markers.

Male production in *D. pulex* can be artificially induced by the addition of methyl farnesoate (MF), a juvenile hormone analog, to the culture medium ^44,46^. To classify MP and NMP phenotypes, approximately 15 adult females (3–4 weeks old) from each clone were exposed to an 800 nM MF working solution for two weeks under long-day photoperiod conditions (14 h light: 10 h dark) at 18°C. MF was diluted in ethanol, stored at –20°C, and added to filtered lake water to reach the working concentration. Individuals were housed singly in 50 mL beakers and transferred and fed algae every two days. During the two-week treatment, each female typically produced 2–4 clutches, allowing assessment of male-production ability. Clones suspected to be NMP were tested in at least two independent replicates. All tested WVA and BRG clones produced 100% female offspring across replicates, confirming their NMP status. Ultimately, we selected five MP clones (NFL87, TEX23, LPB120, POV15, KAP4) and five NMP clones (NFL92, WVA79, KAP106, BRG89, TEX58) for RNA-Seq.

Given that high concentrations of MF may have toxic effects, male induction for RNA-Seq samples was performed using photoperiod manipulation instead. To test responsiveness to photoperiod-induced male production, the five MP clones were reared under both long-day (14L:10D) and short-day (10L:14D) conditions. To reduce maternal effects, first-generation mothers were reared at a density of 10 individuals per 100 mL. From these, 7-day-old females were transferred to establish the second generation. Offspring from the second generation were reared individually and used to assess male-production rates across the first to sixth clutches (Figure S2a). To control for batch effects, each clone was tested in at least two biological replicates (n = 6–10 individuals per replicate).

To get RNA-Seq data, we collected MP clones (NFL87, TEX23, LPB120, POV15, KAP4) and NMP clones (NFL92, WVA79, KAP106, BRG89, TEX58) at five developmental stages: 12h, 36h, and 60h embryos, as well as one-day and seven-day juveniles. To minimize maternal effects, we used the second generation of each clone for sampling (see above). MP female embryos were collected from mothers grown under long-day conditions (18°C), with embryo samples taken at the fourth or fifth clutch (one-month-old), where female production was nearly guaranteed. Male embryos were collected from mothers grown under short-day conditions (18°C), and we sampled from the fourth to sixth clutches based on their peak male-producing rates. NMP embryos were collected from one-month-old mothers under both photoperiod conditions.

All sampled mothers were rinsed with RNase-free water and transferred to three-well porcelain micro spot plates. We briefly immersed the *Daphnia* in DNA/RNA Shield (Zymo Research) for about 30 seconds before dissecting them under a microscope with fine needles (Roboz Surgical RS6062) to separate embryos and mothers. Samples were immediately frozen in liquid nitrogen and stored at −80°C for sequencing.

We collected between 200-545 embryos/juveniles and 18-95 mothers across 70 MP samples [2 photoperiods × (3 embryo stages + 3 mother stages + 2 juvenile stages) × 5 clones = 70] and 70 NMP samples. Total RNA was extracted using either the RNeasy Mini Kit (QIAGEN) or the Direct-zol RNA Miniprep Kit (Zymo Research) and then submitted to Tgen or BGI (the center of Wuhan or Hongkong, China) for rRNA depletion. The quality and integrity of the total RNA were assessed using the NanoDrop spectrophotometer and the 4200 TapeStation (RIN > 8). We designed 155 oligos (Table S4), each at 1 pmol, covering both nuclear and mitochondrial rRNAs of *Daphnia*. Sequencing was performed on the Illumina NovaSeq or DNB-Seq platforms, generating approximately 160Mb of reads per sample.

### Transcriptome analysis

RNA-seq data were cleaned using Trimmomatic to remove adapter sequences and trim low-quality reads. The cleaned reads were then aligned to the KAP4 reference genome using Hisat v2.2.1 ^89^, and unique alignments were extracted. Transcript expression levels (TPM) were estimated using StringTie v2.2.0 ^90^. For each developmental stage, we identified differentially expressed genes (DEGs) by performing a *t*-test, selecting genes with at least a 2-fold difference in average TPM between the five MP and five NMP clones, with a significance threshold of *P* < 0.05. NMP-associated DEGs were defined by comparing MP and NMP clones under short-day photoperiods, but showing no significant differences under long-day conditions.

To identify allele-specific expression, cleaned RNA-seq reads of NMP clones were mapped to the KAP4 reference genome using BWA, allowing alignment of both paired and unpaired reads. Alignments were merged and sorted, and PCR duplicates were removed using Picard with default parameters. Realignment around indels was performed using GATK’s “RealignerTargetCreator” and “IndelRealigner.” Overlapping reads were clipped using BamUtil, and pileup files were generated with Samtools. Allele frequencies across RNA-seq data were estimated using MAPGD. Using NMP markers, we estimated allele-specific expression by calculating the ratio of NMP marker frequencies to the total frequencies within genes.

To identify the *Dfh* co-expression network, we clustered transcripts from DEGs based on their TPM values across MP samples using the Leiden algorithm in Scanpy 1.10 ^92^. The raw TPM values were scaled between 0 and 10, and clustering was performed with 10 neighbors, resolution of 0.5 and minimal distance of 0.05. This analysis yielded nine distinct gene expression clusters, each representing a unique transcriptional pattern in *Daphnia*.

Cluster 5, which included the gene *Dfh*, was selected for further investigation. To explore the gene co-expression network involving *Dfh* and other male-production-related genes within this cluster—particularly those in the juvenile hormone (JH) and ecdysone (20E) pathways, as well as *doublesex*—we reconstructed these pathways by integrating data from KEGG (https://www.genome.jp/kegg/) (“Terpenoid Backbone Biosynthesis” [map00900] and “Insect Hormone Biosynthesis” [map00981]). Using BLAST searches with reference sequences from well-annotated arthropods, we identified 65 genes involved in the JH and 20E pathways in the KAP4 genome, applying thresholds of >80% identity, >50% alignment coverage, and E-value < 10⁻⁶ (Table S2c).

To investigate potential regulatory interactions, we extracted DNA-binding domains of the hormonal responsive genes by searching the conserved domain database ^73^ (CDD) from NCBI. We then analyzed the 1kb upstream region of the *Dfh* transcription start site (TSS) and genic sequences for identifying potential binding sites and predicting interactions using AlphaFold3 ^48^. The predicted structure was visualized using ChimeraX 1.8 ^93^. Additionally, we predicted the signal peptide of *Dfh* to apply SignalP 6.0 ^94^.

### RNA interference and CRISPR

To synthesize the dsRNA template for *Dfh*, *E75*, and *HR3* knock down, we designed by adding the T7 promoter sequence (TAATACGACTCACTATAGGG) to both ends of the dsRNA, which covered most of CDS sequence of targets (Table S4). This sequence was then synthesized by Integrated DNA Technologies and cloned into the pUCIDT-Amp GoldenGate plasmid. We also incorporated restriction enzyme sites: PstI (AACTGCAGAACCAATGCATTGG) at the 5’ end and SpeI (GGACTAGTCC) at the 3’ end of the T7-target-T7 sequence. The resulting plasmid was transformed into 5-alpha Competent *E. coli* cells (High Efficiency, New England Biolabs), and transformed cells were plated on LB agar containing ampicillin (100 mg/mL) for overnight incubation at 37°C to select for resistant colonies.

We selected three single colonies and transferred each to LB broth for overnight culture. After incubation, 1 µL of each culture was used for PCR amplification with the Q5 High-Fidelity 2X Master Mix (New England Biolabs) or the GoTaq(R) Green Master Mix (Promega) using M13 primers. PCR products were purified using the GeneJET PCR Purification Kit (Thermo Fisher Scientific) and sent to the KED Genomics Core at Arizona State University for Sanger sequencing. Correctly sequenced colonies were stored at −80°C.

To prepare for dsRNA synthesis, the pUCIDT-T7-target-T7 plasmids for *Dfh*, *E75*, and *HR3* were recovered from *E. coli* and cultured overnight at 37°C. Plasmids were extracted using the Zyppy Plasmid Miniprep Kit (Zymo Research). The plasmids were then digested with PstI and SpeI (New England Biolabs) for 2 hours at 37°C to ensure complete digestion. The resulting T7-target-T7 template was confirmed by gel electrophoresis, and the correct fragments were recovered using the Zymoclean Gel DNA Recovery Kit (Zymo Research). Using the T7-target-T7 template, dsRNA was synthesized with the HiScribe T7 High Yield RNA Synthesis Kit (New England Biolabs), and purified using the Monarch RNA Cleanup Kit (New England Biolabs).

Three weeks prior to microinjection, we isolated ∼2-week-old female *Daphnia* and maintained them at a density of 10 females per 100 mL in beakers, feeding them every two days with *Scenedesmus obliquus* at a concentration of 150,000 cells/mL. We exposed them to an 800 nM MF working solution for two weeks (under long-day conditions at 18°C). Every 3 days, we transferred the females to new beakers with 800 nM MF to remove any offspring. Sucrose (G-Biosciences) was diluted into filtered COMBO medium to a final concentration of 60 mM.

On the day of microinjection, we selected *Daphnia* exhibiting blue ovaries or black-eyed embryos, indicating the onset of preparing a new reproductive cycle. These individuals were transferred to single wells in 24-well spot plates with adequate COMBO medium, replenished as needed to prevent the wells from drying. We monitored each *Daphnia* for molting and ovulation every five minutes.

After ovulation, we allowed them to rest at room temperature for 10-15 minutes before transferring them to an injection petri dish with minimal COMBO water. The injection dish was prepared by supergluing a glass coverslip (Chemglass Life Sciences; 22×22 mm, 1 ½ thickness) to the lid of a 60×15 mm petri dish (CELLTREAT Scientific Products). A few drops of ice-cold 60 mM sucrose in COMBO were added, and the *Daphnia* were placed on ice for 10-25 minutes, depending on environmental conditions. Under a dissecting microscope (Leica S9i), we separated the embryos from mothers with forceps, minimizing the inclusion of maternal tissue. The embryos were aligned near or against the edge of the coverslip to facilitate injection.

Microneedles (AF100-64-10) were pulled using a P-1000 Glass Puller (Sutter Instrument Company) and beveled with a BV-10 Microelectrode Beveler (Sutter Instrument Company). The final injection pipette specifications were: a shaft length of 5 cm, taper of 5–6 mm, tip diameter of 1–2 µm, and a bevel angle of 30°.We used the FemtoJet 4i Microinjector and InjectMan 4 (Eppendorf) to inject dsRNA (>4 µg/µL) combined with Recombinant Enhanced GFP protein (Abcam) or the GFP protein alone under microscope (Nikon eclipse Ti2). Additional 60 mM sucrose in COMBO was added as needed without disturbing the injected embryos. After one hour, we added 2-3 mL of COMBO to the petri dish and transferred it to a dark room for further incubation.

We engineered potential loss-of-function mutations in the *Dfh* locus using CRISPR–Cas9 genome editing. Two guide RNAs (Table S4) were designed against early exons with the IDT Custom Alt-R™ tool and synthesized by the same provider. Equal molar concentrations of guide RNAs and Cas9 nuclease (IDT) were complexed into ribonucleoproteins (RNPs) in TE buffer, yielding final concentrations of 115 ng/μL for gRNA and 600 ng/μL for Cas9. These RNPs were delivered by microinjection into freshly ovulated embryos from 10–14-day-old females, following a protocol closely paralleling established RNAi injection methods and incorporating modifications outlined by Xu *et al.*^101^. Mutational outcomes at the targeted locus were assayed using the M13(–21) primer genotyping approach (Table S4)^102^, in which PCR fragments were fluorescently labeled and size-resolved on an ABI 3730 analyzer. Fragment lengths were scored with the NCBI OSIRIS platform (https://www.ncbi.nlm.nih.gov/osiris/), allowing efficient detection of Cas9-induced indel variation.

### qPCR validation and morphological assessment

For qPCR validation of RNAi efficiency, total RNA was extracted from ∼15 embryos injected with dsRNA at the 60-hour stage, as well as from embryos injected with only the GFP protein, using the Direct-zol RNA Miniprep Kit. cDNA synthesis was performed with the Luna Universal One-Step RT-qPCR Kit (New England Biolabs). Gene expression levels were quantified using quantitative PCR (qPCR) with the ViiA 7 Real-Time PCR System (Applied Biosystems). Primers for *Dfh*, *E75*, *HR3* and the housekeeping gene alpha-tubulin (served as internal controls) were designed using Primer-BLAST (https://www.ncbi.nlm.nih.gov/tools/primer-blast/). This sequence was then synthesized by Integrated DNA Technologies (Table S4). Relative gene expression was calculated using the ΔΔCt method, with GFP-only injected embryos serving as the control group.

Injected embryos were evaluated for GFP expression on the third day post-injection under a fluorescence microscope (Nikon eclipse Ti2) to confirm successful dsRNA delivery. Moreover, the sex of the injected individuals showing significant GFP expression was determined on the 5th day. For morphology checks, we monitored the ovarian developmental progression of the injected embryos. After 3 weeks, randomly selected embryos that developed into females were assessed using Hematoxylin and Eosin (H&E) staining ^103^. Additionally, uninjected females with the same age were selected as controls for comparative analysis. *Daphnia* were euthanized quickly by transferring them to 70% ethanol and collected into a 15 mL Falcon tube, with excess liquid pipetted off.

The morphological assessment was conducted as follows: Fixation was performed in 10 mL of Bouin’s fixative (Sigma-Aldrich) for 48 hours at room temperature. Following fixation, samples were transferred to 10 mL of 70% ethanol for 24 hours at room temperature. Dehydration, clearing, and paraffin infiltration were done using the HistoCore PEARL Tissue Processor (Leica) on an overnight program. Each Daphnia was embedded in the smallest mold, and double embedding in agarose was utilized when necessary. Sections were then cut at 4–7 µm thickness using the Leica CM1950 Cryostat and stained with H&E (H&E Stain Kit, Abcam) on the Leica Autostainer ST5010 Slide Stainer.

### Single-cell sample preparation, sequencing, and analysis

To minimize maternal effects, we collected second-generation females of KAP4 and KAP106 from a single clone. The second-generation females were maintained under short-day conditions at 18°C. From each group, we sampled over 200 one-month-old females carrying 60-hour embryos, isolating carapace and antennae tissues from both KAP4 and KAP106.

To standardize physiological conditions, *Daphnia* were subjected to a 10-hour starvation period before tissue dissection. We made two washes with autoclaved water at room temperature immediately before dissection. Under a dissecting microscope, carapace and antennae were meticulously excised using fine forceps and micro-scalpels. Tissues were collected in 1X Rinaldini’s solution (800 mg NaCl, 20 mg KCl, 5 mg NaH₂PO₄, 100 mg NaHCO₃, and 100 mg glucose) supplemented with RNase inhibitor (Thermo Fisher Scientific). Samples were centrifuged at 300 × g for 15 minutes at 4°C to pellet the tissues. For tissue dissociation, TrypLE (Gibco) was added to the pellet, and samples were incubated at 37°C, with gentle pipetting every 5 minutes to facilitate enzymatic digestion. The reaction was quenched by adding three volumes of 1X Rinaldini’s solution containing 10% FBS (GenClone). The resulting cell suspension was passed through a 100 µm cell strainer and centrifuged at 500 × g for 15 minutes at 4°C to isolate the cells. The final pellet was resuspended in 150 µL of 1X Rinaldini’s solution, and cell viability was assessed using Trypan blue staining (Invitrogen). Only cell suspensions with a viability exceeding 70% were processed for fixation.

Following the Parse Biosciences Evercode protocol, single cells were fixed using the Evercode Cell Fixation Kit (Parse Biosciences), ensuring RNA integrity was preserved throughout the barcoding and sequencing library preparation stages. This chemical fixation enabled a delay between cell dissociation and library preparation. Using the Evercode WT kit, cells underwent two rounds of split-pooling, during which barcodes were added, assigning each cell a unique combination of cell-specific and molecule-specific indices. After the barcoding, reverse transcription was carried out using barcoded primers to capture mRNA transcripts. The resulting barcoded cDNA was then amplified by PCR to increase yield. Following amplification, the cDNA was fragmented, and sequencing adapters were ligated to the fragments. Library validation included concentration and size distribution assessments using the TapeStation system, confirming typical fragment sizes between 300 and 600 bp. Quantification was performed using Qubit. Finally, the libraries were pooled and sequenced on the Illumina NovaSeq X platform at Tgen, generating approximately 80,000 pair-end reads per cell.

Raw sequencing data were processed using the Parse Biosciences pipeline (https://www.parsebiosciences.com/data-analysis/), which employed the KAP4 reference genome to demultiplex barcoded reads and assign them to individual cells. For each dataset, sampling carapace and antennae tissues from both KAP4 and KAP106, we used Scanpy ^92^ to rigorously filter out low-quality cells and outliers based on the following criteria:

1. A minimum expressed gene count of ≥ 50.
2. The log1p-transformed total count per cell thresholded at 5 median absolute deviations (MADs).
3. The log1p-transformed number of genes with at least one count per cell, thresholded at 5 MADs.
4. The cumulative percentage of counts from the 20 most highly expressed genes per cell, thresholded at 5 MADs.
5. The proportion of mitochondrial reads per cell, thresholded at 3 MADs or >10%.

To further refine cell quality, we removed potential doublets using scDblFinder v1.2 ^94^. Additionally, we retained only genes expressed in at least three cells.

Following cell outlier filtering, we normalized the gene expression matrix to 10,000 reads per cell and log-transformed the data for each tissue (carapace or antennae). Highly variable genes were identified while correcting for batch effects. To mitigate biases in cell attributes, we applied the regression function, adjusting for the total number of counts per cell. Gene expression was then scaled to a maximum of 10. Dimensionality reduction was performed via principal component analysis (PCA), followed by data integration using the Harmony function. Clustering was conducted with the Leiden algorithm (n_neighbors = 20, resolution = 0.6 or 0.8) to define distinct cell populations. Finally, we visualized the clusters using uniform manifold approximation and projection (UMAP), enabling an intuitive exploration of cellular heterogeneity and relationships within the dataset. DEGs between MP and NMP clusters were analyzed using a *t*-test, selecting genes with at least a 2-fold difference in normalized expression between MP and NMP cells within a cluster, with a significance threshold of *P* < 0.05.Cluster annotation was based on *Drosophila* cell markers and a curated gene set involved in cuticle formation, moulting fluid regulation and related functions. We obtained scRNA-seq data of *Drosophila* cells from the FlyCellAtlas ^74^ and selected highly expressed genes in each tissue as potential cell markers. Genes were considered markers if they had read counts > 1000 and were expressed in at least half of the cells. To identify *Drosophila-Daphnia* orthologs, we used OrthoFinder v2.5.5 ^95^, comparing the *Drosophila* and *Daphnia* proteomes obtained from NCBI. Additionally, genes involved in cuticle synthesis and molting fluids were sourced from Muthukrishnan *et al* ^104^.

### Linkage disequilibrium analysis

Due to low sequencing coverage across the 67 NMP clones, direct phasing of *M* and *m* alleles was not feasible. To infer phase, we aligned two haplotypes from each of three high-quality clones (TEX58, KAP106, and NFL92) to the KAP4 reference genome using minimap2, yielding phased genotypes at each locus per clone. From these, we constructed all nine possible *M* × *m* haplotype combinations. Sites that could not be unambiguously phased (e.g., heterozygous *Mm* genotypes such as either be AT or TA) were excluded. We further filtered out genotypes for one given site with low support (allelic count < 3) to eliminate potentially spurious phase calls.

Using this set of phased genotypes derived from six high-confidence haplotypes, we inferred genotypes for the 67 NMP clones. For downstream analyses, we retained only sites where phased genotypes could be confidently inferred from the reference haplotypes and were represented in >30 clones.

LD was quantified using the squared correlation coefficient (r²) ^49^, which quantifies the non-random association of alleles at two biallelic loci. SNPs were categorized into three genomic regions: the inversion region (0–0.6 Mb), the non-inverted portion of the SDR (0.6–1.4 Mb), and the non-SDR (>1.4 Mb) (based on KAP4 coordinates). Within each region, r² was calculated as:

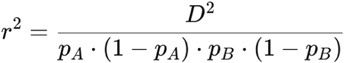

Where *D = f_AB_ – p_A_ · p_B_*. Here, f_AB_ = frequency of the AB haplotype; f_ab_ = frequency of the Ab haplotype; f_aB_ = frequency of the aB haplotype; and f_ab_ = frequency of the ab haplotype. Then, we identify the allele frequencies are p_A_ = f_AB_ + f_Ab_ and p_B_ = f_AB_ + f_aB_.

To validate the existence of a recombination-suppressed region in MP *D. pulex* overlapping the SDR, we analyzed a high-resolution genetic map from a male-producing *D. pulex* clone (KAP4). A total of approximately 15,577 genetic markers from LPB-87 were obtained from Molinier *et al.* ^51^ and mapped to the KAP4 reference genome assembly using BLAST. The corresponding genetic distances for these markers were also taken from the same study. We generated a Marey map by plotting genetic distance (in centiMorgans) against physical position (in megabases) along chromosome I. Regions where genetic distance remained constant or increased only minimally with physical distance were interpreted as indicative of local recombination suppression.

To further explore recombination patterns in related species, we incorporated genetic markers for chromosome I of *D. pulicaria*. The LD profile was derived from Marey maps provided by Wersebe *et al.* ^52^ To identify the homologous SDR region in *D. pulicaria*, whole-genome alignments were performed between the reference *D. pulex* and the *D. pulicaria* LK16 genome assemblies. LD patterns were then extracted from the corresponding genomic regions in *D. pulicaria*.

We collected five *D. obtusa* populations (Table S1d), and analyzed relatedness, which indicates the possibility of clones coming from the same mother. Only the RAP and EBG population displayed low relatedness among clones. LD was calculated based on r² values (as described above) using genotype data from the RAP population. The analysis was restricted to polymorphic sites with fewer than 10 missing genotypes to ensure data quality.

### Origin of gene *Dfh*

To trace the evolutionary origin of the gene *Dfh*, we investigated whether homologs of this gene are present in other *Daphnia* species using GageTracker ^37^, which identifies gene homology through genome pairwise alignments, regardless of annotation completeness in the target species. We obtained genomes of *D. magna* (GCA_020631705.2) and *D. sinensis* (GCA_013167095.2) from GenBank. Additional genomes, including those of *D. arenata*, *D. melanica*, *D. pulicaria* LK6, *D. pulex* BEL2 (a European species), *D. mitsukuri*, *D. catawba*, *D. obtusa*, *D. ambigua*, *D. parvula*, *D. magniceps*, *D. lumholtzi*, and *Simocephalus vetulus*, were sourced from Wei *et al.* and downloaded from iDaphnia database (www.idaphnia.com).

Through comparative analysis, we confirmed the presence of *D*fh homologs across the subgenera *Daphnia*. Notably, incomplete *Dfh-*like sequences were detected in *D. arenata* and *D. catawba*, which we attribute to potential assembly issues rather than true absence of the gene. This widespread conservation across *Daphnia* species suggests an ancient origin of *Dfh* within the genus. To determine the conservation of Dfh protein in sequence and 3D structure, sixteen Dfh protein sequences were aligned using the MUSCLE algorithm in MEGA ^96^ software. AlphaFold3 ^48^ was used to predict the 3D structure of these Dfh proteins and SignalP was used to predict the signal peptides of *Dfh*. The d_N_/d_S_ ratios of *D. pulex* genes were derived from our two previous datasets: one from 10 *D. pulex* populations and another from a population studied over nine years ^72,105^. The final values represent the average of both datasets.

To determine *Dfh* expression in other *Daphnia* species, we tested a single clone each of *D. pulicaria* (MT-20; collected from the U. S., 2023), European *D. pulex* (BEL2; collected from Czech Republic, 2016), *D. mitsukuri* (SZH4; collected from China, 2015), and *D. obtusa* (FS6; collected from the U. S., 2014), all of which have the ability to produce males. Approximately 20 adults, aged three to four weeks, were prepared and exposed to an 800 nM MF solution for two weeks under long-day conditions (18°C). Each individual was housed in a separate 50 mL beaker, transferred to a fresh beaker every two days, and fed algae. Male production was confirmed over the two-week period.

At the end of the two weeks, 60-hour-old embryos were collected, and RNA was extracted using the Direct-zol RNA Miniprep Kit. qPCR was then performed using the Luna Universal One-Step RT-qPCR Kit to assess gene expression levels.

## Supplemental information

**Table S1. Tables related to Figure 1**.

(A) Details of 205 M-specific genes.

(B) Repetitive elements within Chromosome I.

(C) Details of 12 m-specific genes.

(D) Daphnia population or clone information.

(E) Details of unique NMP genotypes

**Table S2. Tables related to Figure 2**.

(A) Summary of RNA-seq data across developmental stages.

(B) DEGs from RNA-seq across developmental stages.

(C) Candidates for NMP genotypes.

(D) Genotype of *Dfh*-KO clones.

(E) Daphnia male-production related genes.

**Table S3. Tables related to Figure 3**.

(A) Top 30 most highly expressed genes in carapce clusters.

(B) Identified gene markers in carapce clusters.

(C) Top 30 most highly expressed genes in antenna clusters.

(D) Identified gene markers in antenna clusters.

(E) DEGs of CC8 between KAP4 and KAP106.

**Table S4. Sequences used in this study.**

---

**Figure S1:**
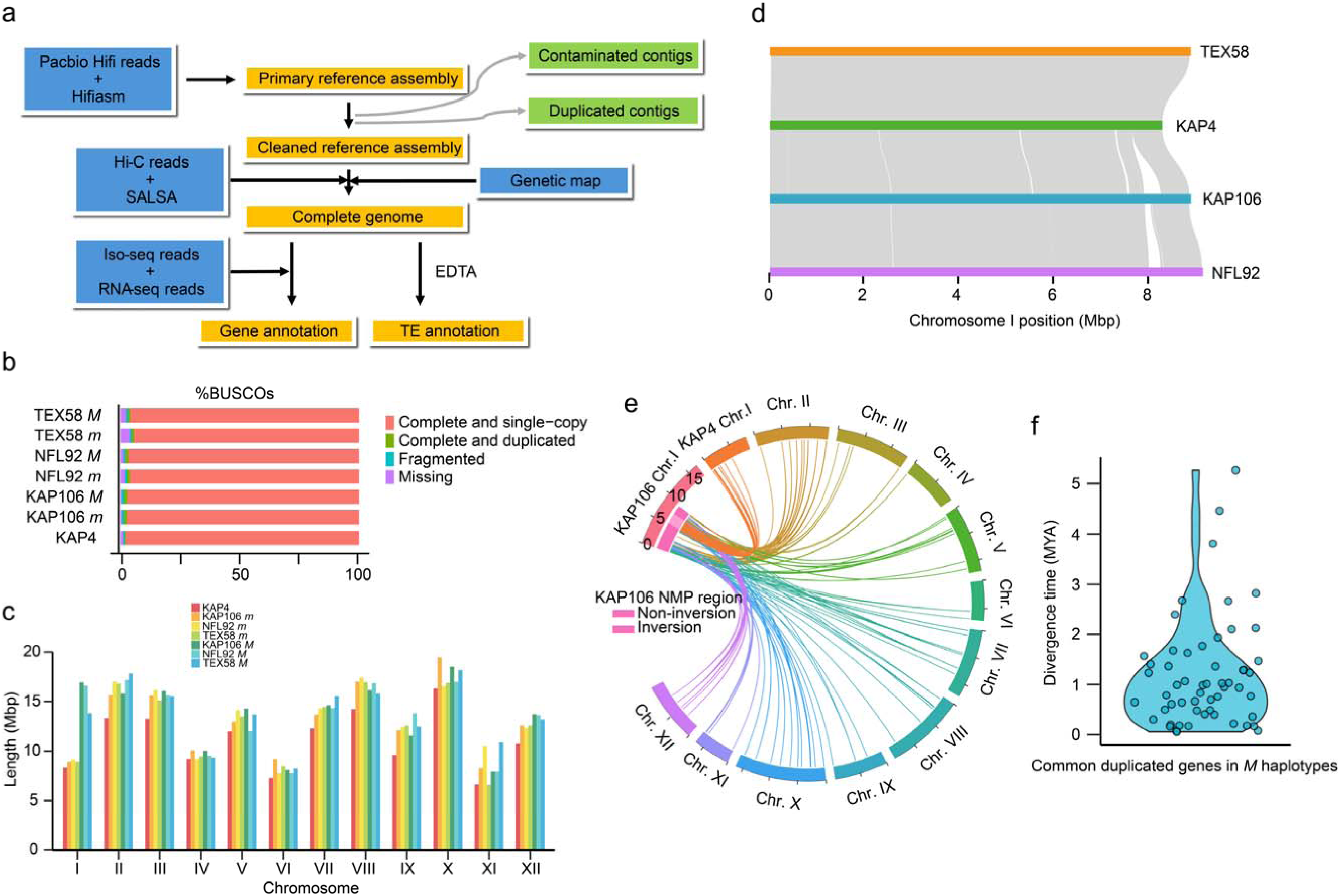
Overview of the genome assembly, related t Figure 1. **a)** Workflow for genome assembly and annotation. **b)** Assessment of genome assembly completeness using BUSCO scores; “M” and “m” indicate assemblies including the *M* and *m* haplotypes, respectively. **c)** Comparison of chromosome lengths across four genomes. **d)** Syntenic comparison of haplotypes from three NMP clones with Chromosome I of the KAP4 reference genome. **e)** Gene synteny between the *M* haplotype of KAP106 and the KAP4 genome. The links highlight homologous genes corresponding to newly duplicated genes in the KAP106 *M* haplotype, indicating the likely genomic positions of their original copies in KAP4. **f)** Distribution of divergence times between newly duplicated genes and their presumed original copies.

**Figure S2:**
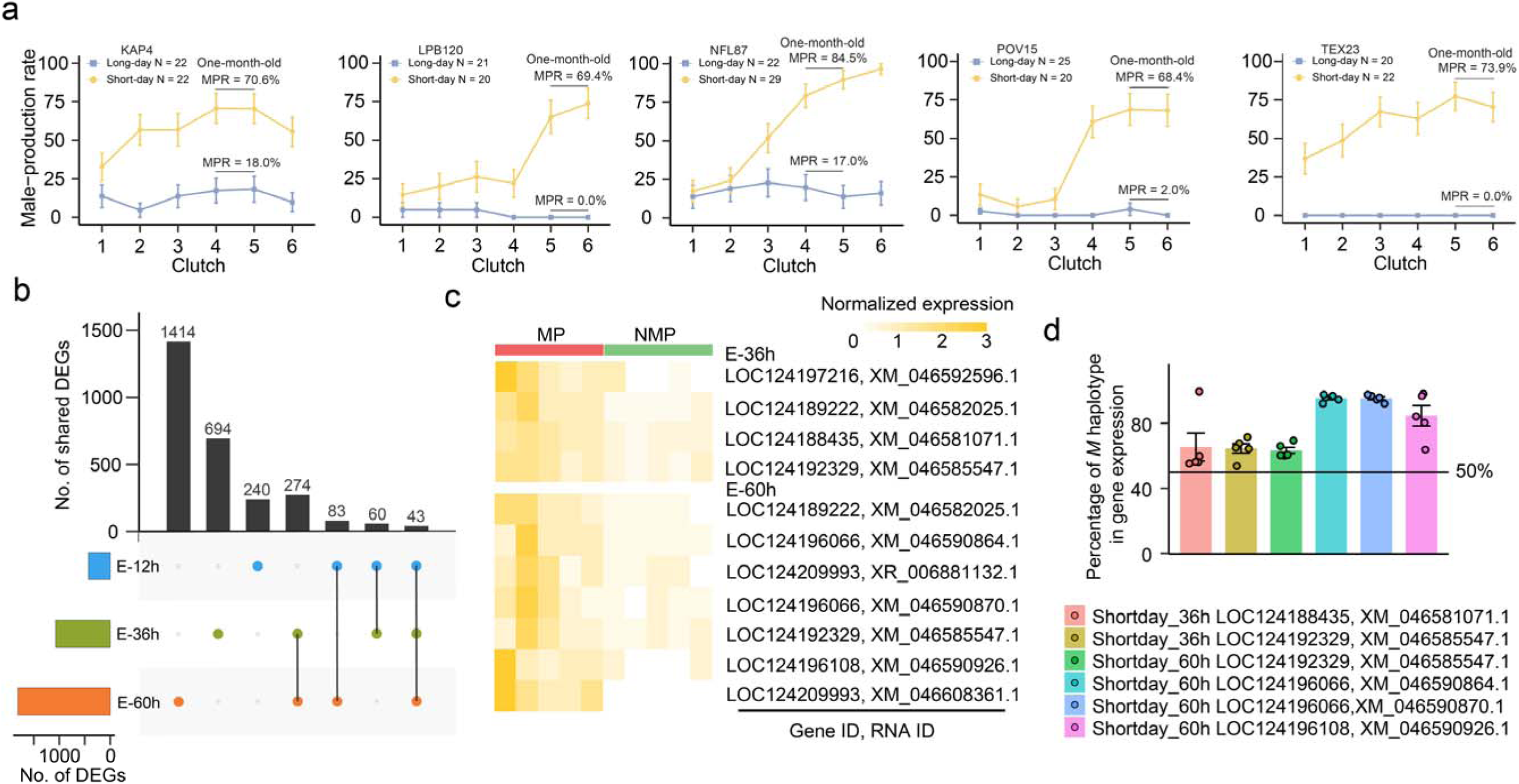
Male-determining gene *Dfh* expression, related to Figure 2. **a)** Comparison of male production rate (MPR) under long-day (LD) and short-day (SD) conditions. Male production rates were assessed in five MP clones (NFL87, TEX23, LPB120, POV15, and KAP4) under both long-day and short-day photoperiods. To minimize maternal effects, first-generation mothers were cultured at a density of 10 individuals per 100 mL. Seven-day-old females from this generation were then transferred to individual 50 mL beakers to produce the second generation. Offspring from this second generation were used to estimate male production rates across the first to sixth clutches. To account for batch effects, two or three independent replicates were conducted per clone (n = 20–29 individuals in total). Data are presented as mean ± S.E.M. for each individual. All clones exhibited peak male production at approximately one month of age. **b)** Upset diagram of DEGs across 3 developmental stages. This diagram illustrates the intersections of DEGs across five developmental stages. DEGs for coding genes were identified by comparing RNA-seq data from five NMP and five MP clones under SD conditions at each stage, excluding DEGs shared under LD conditions. The top panel displays the number of DEGs in each intersection, while the left panel shows the total number of DEGs identified at each developmental stage. Dots in the right panel indicate the presence of specific intersections. **c)** Heatmap of DEGs from the SDR. This heatmap visualizes the expression patterns of DEGs from the SDR. E-12h data are omitted due to no DEGs at that stage. The red and green bars represent five MP and five NMP clones, respectively. Genes were clustered using hierarchical clustering with the “average” linkage method. **d)** Percentage of *M*-haplotype RNA reads for the DEGs listed in **c**. The panel shows the proportion of transcript reads derived from the *M* haplotype relative to total transcript reads. Only values exceeding 50% are shown, indicating reduced expression of the *m* haplotype. Each dot represents an RNA-seq sample; error bars indicate the S.E.M.

**Figure S3:**
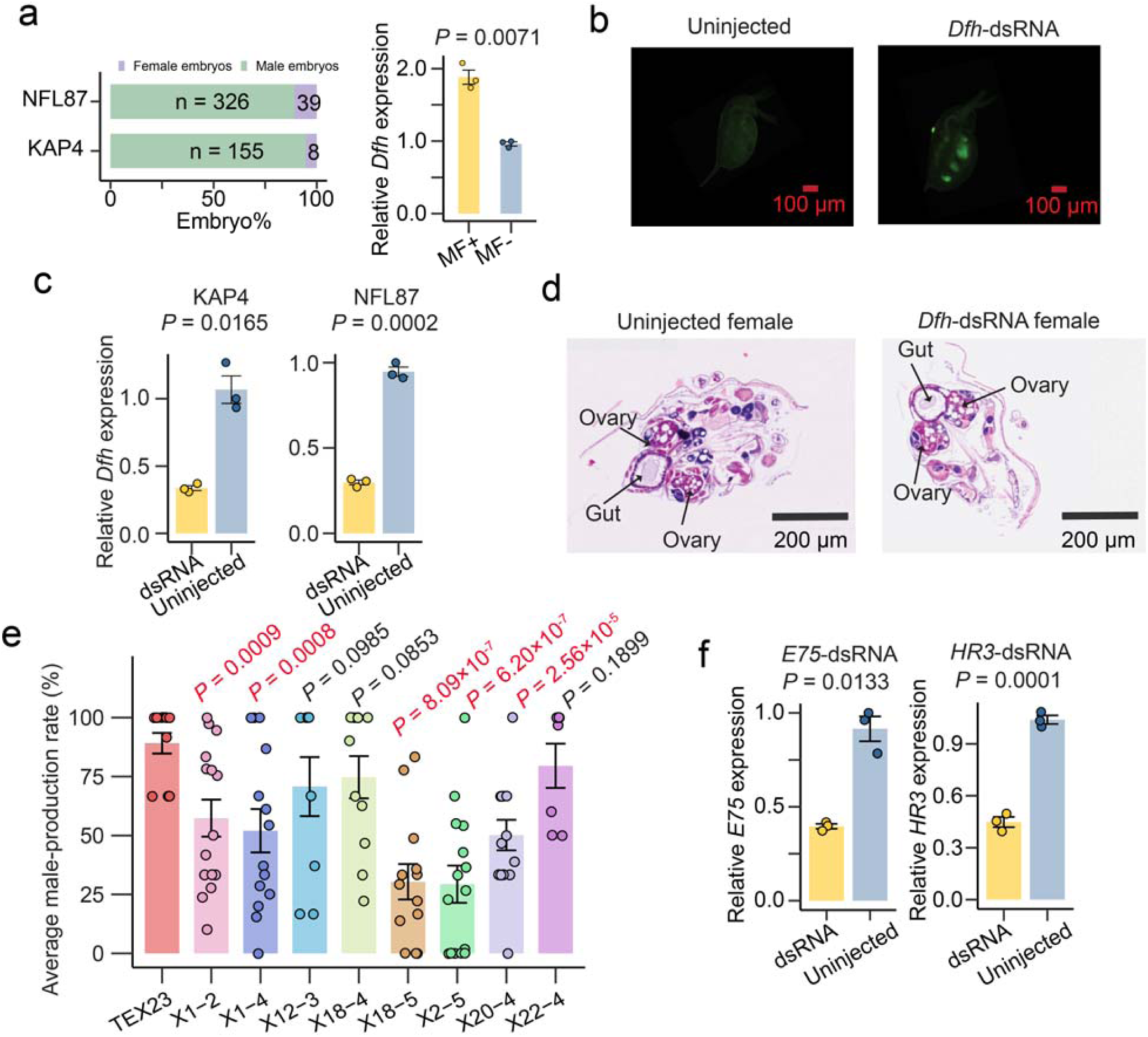
Functional validation of *Dfh*, related to Figure 2. **a)** Effect of methyl farnesoate (MF) on male production in KAP4 and NFL87. MF exposure significantly enhances male production in *Daphnia*. To obtain potential male embryos for microinjection, we assessed male-producing ability under MF treatment. Individuals were exposed to an 800 nM MF solution for two weeks under long-day conditions at 18°C. Under a dissecting microscope, we separated the embryos from mothers with forceps on petri dish, minimizing the inclusion of maternal tissue. The embryos have been transferred to a dark room for further incubation. The number of mothers tested in the KAP4 was 42 and NFL87 was 86. The proportion of sex of developed embryos was estimated (left panel). qPCR validation of *Dfh* Expression comparing the MF-treated (MF+) and untreated group (MF-) (right panel). Over 10 individuals per treatment were collected for RNA extraction, and qPCR analysis was performed. Each dot represents a qPCR replicate, with data presented as mean ± S.E.M. *P*-values were obtained using t-tests comparing MF-treated (MF+) and untreated groups (MF-). **b)** Representative images of *Dfh*-dsRNA-injected and uninjected control *Daphnia* at day three post-injection under fluorescence microscopy. **c)** qPCR validation of *Dfh*-dsRNA knockdown efficiency, measured by *Dfh* expression levels. RNA was extracted from more than 15 individuals at day three post-injection, and qPCR analysis was performed. Each dot represents a qPCR replicate, with data shown as mean ± S.E.M. Statistical significance was assessed using *t*-tests comparing *Dfh*-dsRNA-injected and uninjected groups. **d)** Representative images of hematoxylin-eosin staining of one *Dfh*-dsRNA-injected individual and one uninjected female at weeks three to four post-injection. *Daphnia* were embedded, sectioned, and stained using hematoxylin and eosin procedures. **e)** Average male-production rates for CRISPR knockout lines and the ancestral TEX23 clone under the MF condition. Rates were measured across 7–15 individuals per line over three successive clutches. Statistical differences from TEX23 were evaluated by one-tailed *t*-tests, with significant values (*P* < 0.05) highlighted in red. **f)** qPCR validation of dsRNA knockdown efficiency, measured by *E75/HR3* expression levels. RNA was extracted from more than 15 individuals at day three post-injection, and qPCR analysis was performed.

**Figure S4:**
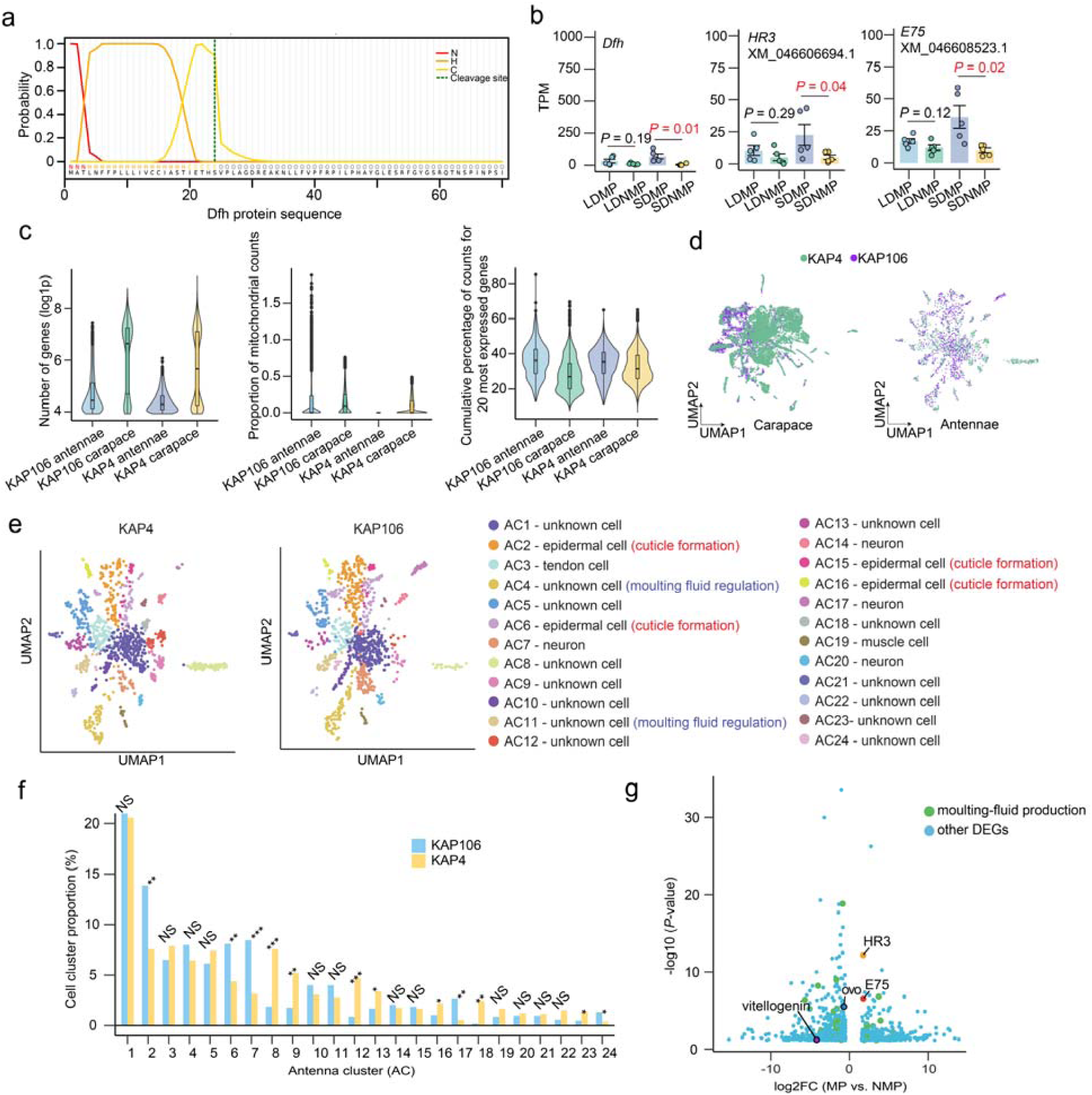
Gene *Dfh* expression in mothers, related to Figure 3. **a)** Prediction of the signal peptide region of Dfh. N, H, and C indicate N-terminal, center hydrophobic and C-terminal region of the signal peptide. **b)** Expression Levels of *Dfh*, *HR3*, and *E75* in MP and NMP mothers. Expression levels of *Dfh*, *HR3*, and *E75* were analyzed in MP and NMP mothers at three developmental stages, characterized by carrying embryos at 12, 36, and 60 hours (M-12h, M-36h, and M-60h stages). Only M-60h data is shown. The TPMvalues are shown for each clone (MP clones: NFL87, TEX23, LPB120, POV15, and KAP4 and NMP clones: NFL92, WVA79, KAP106, BRG89, and TEX58). Each dot represents a clone, with data presented as mean ± S.E.M. *P*-values were calculated using *t*-tests comparing MP and NMP clones under LD and SD conditions, with significant values (*P* < 0.05) highlighted in red. For *HR3* and *E75*, only transcripts co-expressed with *Dfh* in cluster 5 (as shown in Figure 2g) are displayed. **c)** Violin plots displaying the distribution of the number of detected genes, the proportion of mitochondrial read counts relative to the total, and the cumulative percentage of counts contributed by the 20 most highly expressed genes, for four scRNA-seq datasets. **d)** Comparison of KAP4 and KAP106 cells in carapace and antennae. Cells from both samples are visualized as colored dots on the plot. **e)** Cell clustering. Cells were clustered using the UMAP algorithm based on gene expression profiles, identifying 24 distinct cell clusters in the antennae (represented by colored dots). Cluster annotation was based on *Drosophila* cell markers and a curated gene set involved in cuticle formation (underlined), moulting fluid regulation (black border), and other related functions. **f)** Proportion of antenna cell clusters. A chi-square test was used to compare the proportion of each antenna cell cluster between KAP4 and KAP106 relative to the total cell count. Statistical significance is indicated as follows: NS (not significant), * (0.001 ≤ *P* < 0.05), ** (1e-6 ≤ *P* < 0.001), *** (*P* < 1e-6). **g)** volcano plot of DEGs between KAP4 and KAP4-CC8 cells.

**Figure S5:**
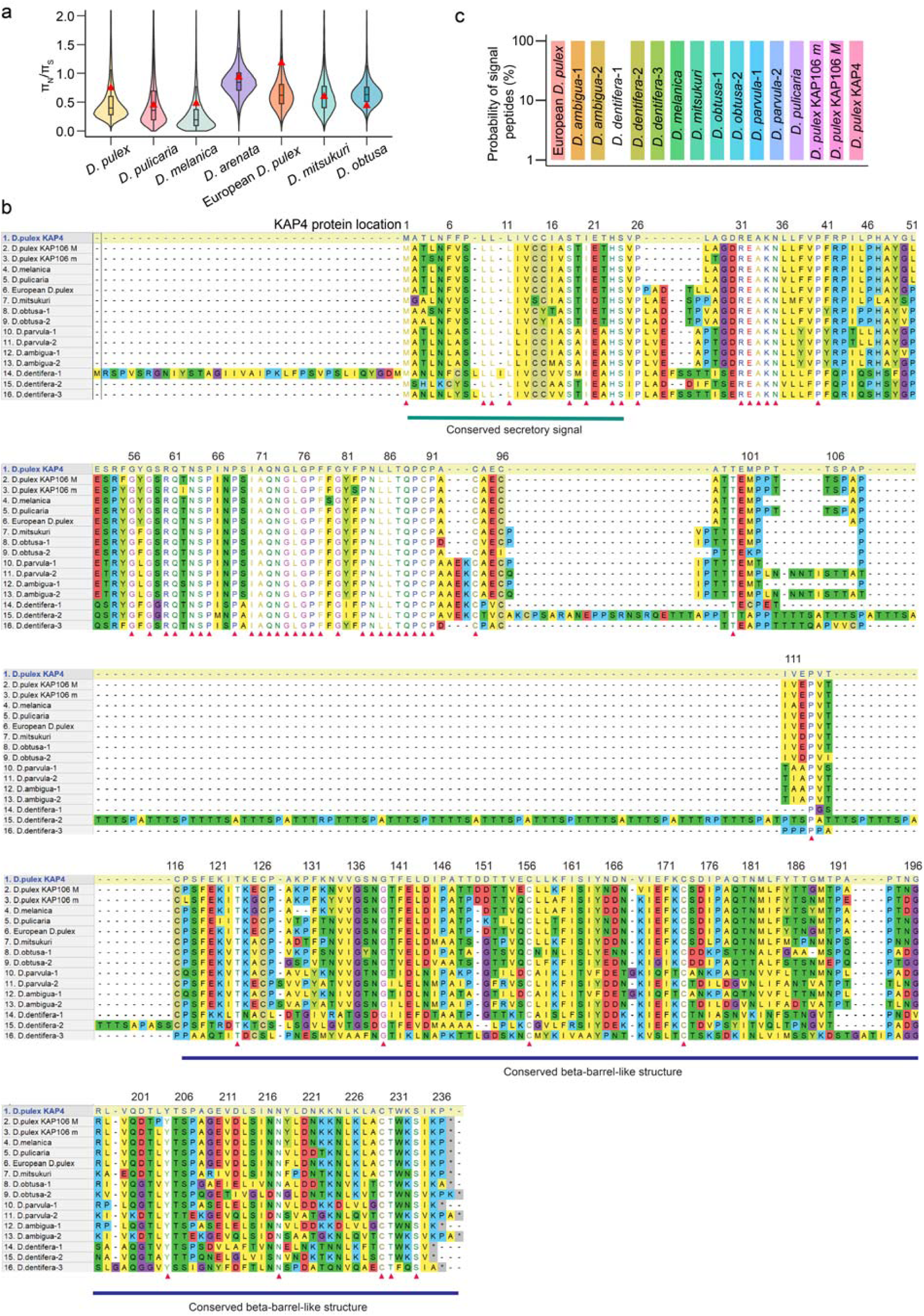
*Dfh* paralogs, related to Figure 5. **a)** Violin plots show the distribution of π_N_/π_S_ ratios for genes across *Daphnia* species. Ratios were estimated within populations and then averaged across populations for each gene. Analyses were restricted to KAP4 genes and their homologs. Red triangles indicate ratios of *Dfh* genes. **b)** Alignment of Dfh proteins across *Daphnia* species. Sixteen Dfh protein sequences were aligned using the MUSCLE algorithm in MEGA software. Amino acids are color-coded based on their biochemical properties: yellow (A, M, F, I, V, L), olive (C), green (N, Q, S, T, W), aqua (D, E), blue (P), red (R, K), fuchsia (G), teal (H), and lime (Y). Red triangles indicate sites conserved across all species, and the blue line highlights the beta-barrel-like domain. **c)** Predicted probabilities of secretory signal across *Daphnia* species.

**Figure S6:**
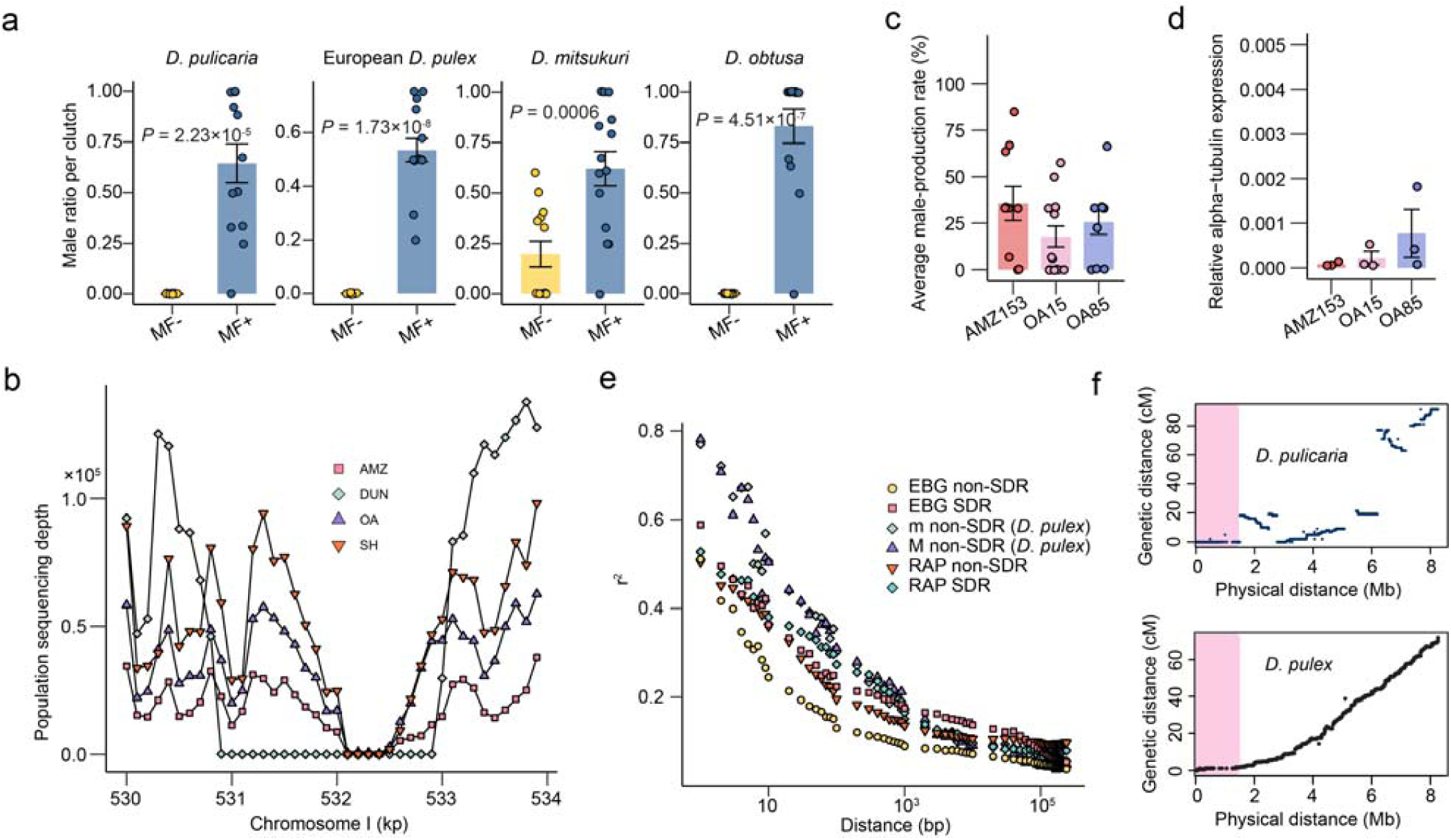
Origin and Evolution of *Dfh*, related to Figure 5. **a)** Male production capacity across four *Daphnia* species. One clone each of *D. pulicaria*, European *D. pulex*, *D. mitsukuri*, and *D. obtusa* was exposed to MF for two weeks under long-day conditions at 18°C. Sample sizes for MF-treated groups (MF+) were 13, 14, 14, and 13 individuals, respectively; untreated groups (MF−) included 13, 12, 13, and 12 individuals. Each dot represents a single individual. Data are shown as mean ± S.E.M. of the average male proportion over the two-week period. *P*-values were derived from *t*-tests comparing MF+ and MF− groups. **b)** Sequencing depth across four *D. arenata* populations. Overall population coverage is indicated by Y-axis. Nearly the entire β-barrel–like domain (532071–532679) shows zero read coverage across all *D. arenata* clones, consistent with complete loss of this region. **c)** Average male-production rates for three representative *D. arenata* clones under the MF condition. Rates were measured across 10–14 individuals per clone over three successive clutches. **d)** *Dfh* expression in three representative *D. arenata* clones under the MF condition. Individuals were exposed to MF for at least two weeks, and 60-hour embryos from >15 individuals per clone were collected for RNA extraction. Expression was quantified by qPCR as mean ± S.E.M. from three primer sets spanning exons 1– 2, 2–3, and 3–4, with each value representing the average of three technical replicates. **e)** Linkage disequilibrium in *D. obtusa* populations EBG and RAP. LD, quantified by the squared correlation coefficient (r²), was calculated from population genomic data of EBG and RAP. **f)** Genetic map of a male-producing *D. pulicaria* and *D. pulex*. The Marey map was constructed using genetic markers, with marker positions aligned to the *D. pulicaria* LK16 or *D. pulex* KAP4 reference genome, respectively. The pink highlights a homologous region of *D. pulex* SDR.

## References

1. Charlesworth, B. & Charlesworth, D. Elements of evolutionary genetics, (Roberts and Co. Publishers, Greenwood Village, Colo, 2010).

2. Charnov, E.L. The theory of sex allocation, (Princeton university press, 2020).

3. Bell, G. The masterpiece of nature: the evolution and genetics of sexuality, (Routledge, 2019).

4. Van Doorn, G.S. Patterns and mechanisms of evolutionary transitions between genetic sex-determining systems. Cold Spring Harbor perspectives in biology 6, a017681 (2014).

5. Straková, B., Rovatsos, M., Kubička, L. & Kratochvíl, L. Evolution of sex determination in amniotes: did stress and sequential hermaphroditism produce environmental determination? Bioessays 42, e2000050 (2020).

6. Zhu, Z., Younas, L. & Zhou, Q. Evolution and regulation of animal sex chromosomes. Nat Rev Genet 26, 59–74 (2025).

7. Goyal, R., Baxter, E. & Van Doren, M. sisterless A is required for activation of Sex lethal in the Drosophila germline. (eLife Sciences Publications, Ltd, 2024).

8. Beukeboom, L.W. & Perrin, N. The evolution of sex determination, (Oxford University Press, USA, 2014).

9. Charlesworth, D., Charlesworth, B. & Marais, G. Steps in the evolution of heteromorphic sex chromosomes. Heredity 95, 118–128 (2005).

10. Cline, T.W. Evidence that sisterless-a and sisterless-b are two of several discrete “numerator elements” of the X/A sex determination signal in *Drosophila* that switch Sxl between two alternative stable expression states. Genetics 119, 829–62 (1988).

11. Katsura, Y., Kondo, H.X., Ryan, J., Harley, V. & Satta, Y. The evolutionary process of mammalian sex determination genes focusing on marsupial SRYs. BMC Evol Biol 18, 3 (2018).

12. Sinclair, A.H. et al. A gene from the human sex-determining region encodes a protein with homology to a conserved DNA-binding motif. Nature 346, 240–4 (1990).

13. Koopman, P., Gubbay, J., Vivian, N., Goodfellow, P. & Lovell-Badge, R. Male development of chromosomally female mice transgenic for Sry. Nature 351, 117–21 (1991).

14. Sardell, J.M., Josephson, M.P., Dalziel, A.C., Peichel, C.L. & Kirkpatrick, M. Heterogeneous histories of recombination suppression on stickleback sex chromosomes. Mol Biol Evol 38, 4403–4418 (2021).

15. Hattori, R.S. et al. The duplicated y-specific amhy gene is conserved and linked to maleness in silversides of the genus odontesthes. Genes (Basel) 10(2019).

16. Koyama, T. et al. A SNP in a steroidogenic enzyme is associated with phenotypic sex in seriola fishes. Curr Biol 29, 1901–1909.e8 (2019).

17. Chikami, Y., Okuno, M., Toyoda, A., Itoh, T. & Niimi, T. Evolutionary history of sexual differentiation mechanism in insects. Mol Biol Evol 39(2022).

18. Verhulst, E.C., van de Zande, L. & Beukeboom, L.W. Insect sex determination: it all evolves around transformer. Curr Opin Genet Dev 20, 376–83 (2010).

19. Koopman, P. Sex determination: the power of DMRT1. Trends Genet 25, 479–81 (2009).

20. Wright, A.E., Dean, R., Zimmer, F. & Mank, J.E. How to make a sex chromosome. Nature communications 7, 12087 (2016).

21. Charlesworth, B. & Charlesworth, D. A model for the evolution of dioecy and gynodioecy. The American Naturalist 112, 975–997 (1978).

22. van Doorn, G.S. & Kirkpatrick, M. Turnover of sex chromosomes induced by sexual conflict. Nature 449, 909–12 (2007).

23. Rice, W.R. The accumulation of sexually antagonistic genes as a selective agent promoting the evolution of reduced recombination between primitive sex chromosomes. Evolution 41, 911–914 (1987).

24. Lenormand, T. & Roze, D. Y recombination arrest and degeneration in the absence of sexual dimorphism. Science 375, 663–666 (2022).

25. Abdel-Haleem, H. The origins of genome architecture. Journal of Heredity 98, 633–634 (2007).

26. Jeffries, D.L., Gerchen, J.F., Scharmann, M. & Pannell, J.R. A neutral model for the loss of recombination on sex chromosomes. Philosophical Transactions of the Royal Society B: Biological Sciences 376, 20200096 (2021).

27. Charlesworth, B., Coyne, J.A. & Barton, N.H. The relative rates of evolution of sex chromosomes and autosomes. The American Naturalist 130, 113–146 (1987).

28. Lenormand, T., Fyon, F., Sun, E. & Roze, D. Sex chromosome degeneration by regulatory evolution. Curr Biol 30, 3001–3006.e5 (2020).

29. Charlesworth, D. The timing of genetic degeneration of sex chromosomes. Philos Trans R Soc Lond B Biol Sci 376, 20200093 (2021).

30. Ebert, D. Daphnia as a versatile model system in ecology and evolution. Evodevo 13, 16 (2022).

31. Peterson, J.K., Kashian, D.R. & Dodson, S.I. Methoprene and 20-OH-ecdysone affect male production in Daphnia pulex. Environ Toxicol Chem 20, 582–8 (2001).

32. Charlesworth, B. Model for evolution of Y chromosomes and dosage compensation. Proceedings of the National Academy of Sciences 75, 5618–5622 (1978).

33. Kato, Y., Kobayashi, K., Watanabe, H. & Iguchi, T. Environmental sex determination in the branchiopod crustacean *Daphnia magna*: deep conservation of a Doublesex gene in the sex-determining pathway. PLoS Genet 7, e1001345 (2011).

34. Ye, Z., Molinier, C., Zhao, C., Haag, C.R. & Lynch, M. Genetic control of male production in Daphnia pulex. Proc Natl Acad Sci U S A 116, 15602–15609 (2019).

35. Pannell, J.R. & Jordan, C.Y. Evolutionary transitions between hermaphroditism and dioecy in animals and plants. Annual Review of Ecology, Evolution, and Systematics 53, 183–201 (2022).

36. Abbott, J.K., Nordén, A.K. & Hansson, B. Sex chromosome evolution: historical insights and future perspectives. Proceedings of the Royal Society B: Biological Sciences 284, 20162806 (2017).

37. Jordão, R. et al. Mechanisms of action of compounds that enhance storage lipid accumulation in *Daphnia magna*. Environ Sci Technol 50, 13565–13573 (2016).

38. Ye, Z. et al. A new reference genome assembly for the microcrustacean *Daphnia pulex*. G3: Genes, Genomes, Genetics 7, 1405–1416 (2017).

39. Keith, N. et al. High mutational rates of large-scale duplication and deletion in *Daphnia pulex*. Genome Res 26, 60–9 (2016).

40. Hannas, B.R. & LeBlanc, G.A. Expression and ecdysteroid responsiveness of the nuclear receptors HR3 and E75 in the crustacean *Daphnia magna*. Mol Cell Endocrinol 315, 208–18 (2010).

41. Molinier, C. et al. Evolution of gene expression during a transition from environmental to genetic sex determination. Mol Biol Evol 36, 1551–1564 (2019).

42. Mathers, T.C. et al. Transition in sexual system and sex chromosome evolution in the tadpole shrimp *Triops cancriformis*. Heredity (Edinb) 115, 37–46 (2015).

43. Hebert, P.D.N. The population bilogy of Daphnia (crustacea, daphnidae). Biological Reviews 53, 387–426 (1978).

44. Toyota, K. et al. Methyl farnesoate synthesis is necessary for the environmental sex determination in the water flea Daphnia pulex. J Insect Physiol 80, 22–30 (2015).

45. Olmstead, A.W. & Leblanc, G.A. Juvenoid hormone methyl farnesoate is a sex determinant in the crustacean Daphnia magna. J Exp Zool 293, 736–9 (2002).

46. Toyota, K. et al. NMDA receptor activation upstream of methyl farnesoate signaling for short day-induced male offspring production in the water flea, *Daphnia pulex*. BMC Genomics 16, 186 (2015).

47. Traag, V.A., Waltman, L. & van Eck, N.J. From Louvain to Leiden: guaranteeing well-connected communities. Scientific Reports 9, 5233 (2019).

48. Abramson, J. et al. Accurate structure prediction of biomolecular interactions with AlphaFold 3. Nature 630, 493–500 (2024).

49. Lewontin, R.C. The interaction of selection and linkage. I. General considerations; Heterotic models. Genetics 49, 49–67 (1964).

50. Cornetti, L., Fields, P.D., Van Damme, K. & Ebert, D. A fossil-calibrated phylogenomic analysis of Daphnia and the Daphniidae. Mol Phylogenet Evol 137, 250–262 (2019).

51. Molinier, C., Lenormand, T. & Haag, C.R. No recombination suppression in asexually produced males of Daphnia pulex. Evolution 77, 1987–1999 (2023).

52. Wersebe, M.J., Sherman, R.E., Jeyasingh, P.D. & Weider, L.J. The roles of recombination and selection in shaping genomic divergence in an incipient ecological species complex. Mol Ecol 32, 1478–1496 (2023).

53. Charlesworth, D., Charlesworth, B. & Marais, G. Steps in the evolution of heteromorphic sex chromosomes. Heredity (Edinb) 95, 118–28 (2005).

54. Wright, A.E., Dean, R., Zimmer, F. & Mank, J.E. How to make a sex chromosome. Nat Commun 7, 12087 (2016).

55. Soh, Y.Q. et al. Sequencing the mouse Y chromosome reveals convergent gene acquisition and amplification on both sex chromosomes. Cell 159, 800–13 (2014).

56. Bergero, R. & Charlesworth, D. The evolution of restricted recombination in sex chromosomes. Trends Ecol Evol 24, 94–102 (2009).

57. Natri, H., Garcia, A.R., Buetow, K.H., Trumble, B.C. & Wilson, M.A. The pregnancy pickle: evolved immune compensation due to pregnancy underlies sex differences in human diseases. Trends Genet 35, 478–488 (2019).

58. Kondo, M. et al. Genomic organization of the sex-determining and adjacent regions of the sex chromosomes of medaka. Genome Res 16, 815–26 (2006).

59. Kikuchi, K. & Hamaguchi, S. Novel sex-determining genes in fish and sex chromosome evolution. Developmental Dynamics 242, 339–353 (2013).

60. Kortum, R.L. & Lewis, R.E. The molecular scaffold *KSR1* regulates the proliferative and oncogenic potential of cells. Mol Cell Biol 24, 4407–16 (2004).

61. Nguyen, A. et al. Kinase suppressor of Ras (*KSR*) is a scaffold which facilitates mitogen-activated protein kinase activation in vivo. Mol Cell Biol 22, 3035–45 (2002).

62. Whitmarsh, A.J. Regulation of gene transcription by mitogen-activated protein kinase signaling pathways. Biochim Biophys Acta 1773, 1285–98 (2007).

63. Hennig, J. et al. Structural basis for the assembly of the *Sxl-Unr* translation regulatory complex. Nature 515, 287–90 (2014).

64. Moschall, R., Strauss, D., García-Beyaert, M., Gebauer, F. & Medenbach, J. *Drosophila* Sister-of-Sex-lethal is a repressor of translation. Rna 24, 149–158 (2018).

65. Reisser, C.M. et al. Transition from environmental to partial genetic sex determination in Daphnia through the evolution of a female-determining incipient W chromosome. Mol Biol Evol 34, 575–588 (2017).

66. Deng, H.W. & Lynch, M. Inbreeding depression and inferred deleterious-mutation parameters in Daphnia. Genetics 147, 147–55 (1997).

67. Haag, C.R., Hottinger, J.W., Riek, M. & Ebert, D. Strong inbreeding depression in a Daphnia metapopulation. Evolution 56, 518–26 (2002).

68. Haag, C.R., Hotinger, J.W., Riex, M. & Ebert, D. Strong inbreeding depression in a Daphnia metapopulation. Evolution 56, 518–526 (2002).

69. Deng, H.-W. & Lynch, M. Inbreeding depression and inferred deleterious-mutation parameters in Daphnia. Genetics 147, 147–155 (1997).

70. Ye, Z., Pfrender, M.E. & Lynch, M. Evolutionary genomics of sister species differing in effective population sizes and recombination rates. Genome Biol Evol 15(2023).

71. Maruki, T., Ye, Z. & Lynch, M. Evolutionary genomics of a subdivided species. Mol Biol Evol 39(2022).

72. Lynch, M., Wei, W., Ye, Z. & Pfrender, M. The genome-wide signature of short-term temporal selection. Proc Natl Acad Sci U S A 121, e2307107121 (2024).

73. Wang, J. et al. The conserved domain database in 2023. Nucleic Acids Res 51, D384–d388 (2023).

74. Li, H. et al. Fly Cell Atlas: A single-nucleus transcriptomic atlas of the adult fruit fly. Science 375, eabk2432 (2022).

75. Cheng, H. et al. Haplotype-resolved assembly of diploid genomes without parental data. Nat Biotechnol 40, 1332–1335 (2022).

76. Camacho, C. et al. BLAST+: architecture and applications. BMC Bioinformatics 10, 421 (2009).

77. Guan, D. et al. Identifying and removing haplotypic duplication in primary genome assemblies. Bioinformatics 36, 2896–2898 (2020).

78. Ghurye, J. et al. Integrating Hi-C links with assembly graphs for chromosome-scale assembly. PLoS Comput Biol 15, e1007273 (2019).

79. Durand, N.C. et al. Juicer provides a one-click system for analyzing loop-resolution hi-c experiments. Cell Syst 3, 95–8 (2016).

80. Manni, M., Berkeley, M.R., Seppey, M., Simão, F.A. & Zdobnov, E.M. BUSCO update: novel and streamlined workflows along with broader and deeper phylogenetic coverage for scoring of eukaryotic, prokaryotic, and viral genomes. Mol Biol Evol 38, 4647–4654 (2021).

81. Li, H. Minimap2: pairwise alignment for nucleotide sequences. Bioinformatics 34, 3094–3100 (2018).

82. Bolger, A.M., Lohse, M. & Usadel, B. Trimmomatic: a flexible trimmer for Illumina sequence data. Bioinformatics 30, 2114–20 (2014).

83. Li, H. & Durbin, R. Fast and accurate short read alignment with Burrows-Wheeler transform. Bioinformatics 25, 1754–60 (2009).

84. McKenna, A. et al. The Genome Analysis Toolkit: a MapReduce framework for analyzing next-generation DNA sequencing data. Genome Res 20, 1297–303 (2010).

85. Jun, G., Wing, M.K., Abecasis, G.R. & Kang, H.M. An efficient and scalable analysis framework for variant extraction and refinement from population-scale DNA sequence data. Genome Res 25, 918–25 (2015).

86. Danecek, P. et al. Twelve years of SAMtools and BCFtools. Gigascience 10(2021).

87. Maruki, T. & Lynch, M. Genotype-frequency estimation from high-throughput sequencing data. Genetics 201, 473–86 (2015).

88. Goel, M. & Schneeberger, K. plotsr: visualizing structural similarities and rearrangements between multiple genomes. Bioinformatics 38, 2922–2926 (2022).

89. Kim, D., Paggi, J.M., Park, C., Bennett, C. & Salzberg, S.L. Graph-based genome alignment and genotyping with HISAT2 and HISAT-genotype. Nature Biotechnology 37, 907–915 (2019).

90. Kovaka, S. et al. Transcriptome assembly from long-read RNA-seq alignments with StringTie2. Genome Biol 20, 278 (2019).

91. Teufel, F. et al. SignalP 6.0 predicts all five types of signal peptides using protein language models. Nature Biotechnology 40, 1023–1025 (2022).

92. Wolf, F.A., Angerer, P. & Theis, F.J. SCANPY: large-scale single-cell gene expression data analysis. Genome Biol 19, 15 (2018).

93. Meng, E.C. et al. UCSF ChimeraX: Tools for structure building and analysis. Protein Sci 32, e4792 (2023).

94. Germain, P.L., Lun, A., Garcia Meixide, C., Macnair, W. & Robinson, M.D. Doublet identification in single-cell sequencing data using scDblFinder. F1000Res 10, 979 (2021).

95. Emms, D.M. & Kelly, S. OrthoFinder: solving fundamental biases in whole genome comparisons dramatically improves orthogroup inference accuracy. Genome Biology 16, 157 (2015).

96. Kumar, S. et al. MEGA12: molecular evolutionary genetic analysis version 12 for adaptive and green computing. Molecular Biology and Evolution 41, msae263 (2024).

97. Ou, S. et al. Differences in activity and stability drive transposable element variation in tropical and temperate maize. Genome Res 34, 1140–1153 (2024).

98. Kilham, S.S., Kreeger, D.A., Lynn, S.G., Goulden, C.E. & Herrera, L. COMBO: a defined freshwater culture medium for algae and zooplankton. Hydrobiologia 377, 147–159 (1998).

99. Gabriel, L. et al. BRAKER3: Fully automated genome annotation using RNA-seq and protein evidence with GeneMark-ETP, AUGUSTUS, and TSEBRA. Genome Res 34, 769–777 (2024).

100. Ou, S. et al. Benchmarking transposable element annotation methods for creation of a streamlined, comprehensive pipeline. Genome Biol 20, 275 (2019).

101. Xu, S. et al. The mosaicism of Cas-induced mutations and pleiotropic effects of scarlet gene in an emerging model system. Heredity 134, 221–233 (2025).

102. Schuelke, M. An economic method for the fluorescent labeling of PCR fragments. Nature Biotechnology 18, 233–234 (2000).

103. Ngu, M.S. et al. A web-based histology atlas for the freshwater sentinel species *Daphnia magna*. Science of The Total Environment 958, 177930 (2025).

104. Muthukrishnan, S. et al. Chapter One - Chitin in insect cuticle. in Advances in Insect Physiology, Vol. 62 (ed. Sugumaran, M.) 1-110 (Academic Press, 2022).

105. Ye, Z., Wei, W., Pfrender, M.E. & Lynch, M. Evolutionary insights from a large-scale survey of population-genomic variation. Mol Biol Evol 40(2023).

